# Multiple paths lead to salt tolerance - pre-adaptation vs dynamic responses from two closely related extremophytes

**DOI:** 10.1101/2021.10.23.465591

**Authors:** Kieu-Nga Tran, Guannan Wang, Dong-Ha Oh, John C. Larkin, Aaron P. Smith, Maheshi Dassanayake

## Abstract

Salt tolerance is a complex trait with much of the underlying genetic variation and integrated response strategies yet to be discovered from stress adapted plants. *Schrenkiella parvula* and *Eutrema salsugineum* are salt-tolerant extremophytes related to *Arabidopsis thaliana* in Brassicaceae. We investigated their response strategies contrasted against the salt-sensitive model, *A. thaliana* to cope with salt stresses via transcriptomic, metabolomic, and ionomic adjustments. The extremophytes exemplified divergent routes to achieve nutrient balance, build osmotolerance, boost antioxidant capacity, and extend transcriptomic support for modified ion transport and stress signaling. Those led to similar molecular phenotypes adapted to salt stress in the extremophytes, absent in *A. thaliana*. The predominant transcriptomic signals in all three species were associated with salt stress. However, root architecture modulation mediated by negative regulation of auxin and ABA signaling supported minimally-affected root growth unique to each extremophyte during salt treatments. Overall, *E. salsugineum* exhibited pre-adapted responses at the metabolome level, whereas *S. parvula* showed dynamic metabolomic responses coupled to a pre-adapted transcriptome to survive salt stress. Our work shows that the two extremophytes share common salt tolerance features, but differ substantially in pathways leading to the convergent, adaptive traits.

## Introduction

Plants differ greatly in their tolerance to salt stress and there is a metabolic cost for adaptation to salt stress, reflected by differences in growth and yield when grown in high saline soils ^1, 2^. Approximately 2% of angiosperms are adapted to grow in high saline environments while the remaining 98%, including most crop plants, are highly sensitive to salt stress ^3, 4^. Previous studies have largely used salt-sensitive model plants or crops to understand genetic mechanisms underlying salt tolerance. Consequently, we understand more about functional aspects of individual genes or more specifically their maladapted variants and how they function in genomic backgrounds sensitive to salinity stress. However, insights from comparative studies using salt-tolerant extremophytes contrasted with a salt-sensitive model offer a promising prospect for discovery of evolutionarily successful tolerance mechanisms that are absent or underdeveloped in the salt-sensitive plants ^5, 6^. This allows us to rethink the functional space of genes and their influence on co-regulated pathways leading to salt tolerance unexplored before.

Excess salt exerts osmotic, oxidative, ionic, and water-deficit stresses ^7–9^. Multiple genetic mechanisms mediated by ABA-dependent as well as ABA-independent pathways have been shown to modulate salt stress responses in *A. thaliana*, all major crops, and selected halophytes ^8, 10, 11, 108^. These studies collectively support the view that salt stress-adapted plants show a highly coordinated response to survive salt stress that requires synergistic coordination between root and shoot responses. Despite the complexity of adapting to salt stress at the whole plant level, angiosperm lineages have evolved this complex trait repeatedly ^12, 13^. Therefore, it is one of the main complex traits studied for convergent evolution in plants. Despite the significant body of work on salt stress adaptation in plants, we still have a large gap in understanding which pathways synergistically act to achieve salt stress adaptation, and how a common set of orthologs from closely related plants mediate this adaptation ^14^.

*Schrenkiella parvula* and *Eutrema salsugineum* (Brassicaceae) are two leading extremophyte models equipped with foundational genomic resources which facilitate comparative studies with *Arabidopsis thaliana* ^6, 15, 16^. These extremophytes show remarkable salt resilient growth even at salinities reaching seawater strengths ^6, 17, 18^. While *S. parvula* is found near salt lakes in the Irano-Turanian region ^4, 19^, *E. salsugineum* has a wider distribution from coastal to inland saline fields in the northern temperate to sub-arctic regions ^20^. Despite several studies highlighting metabolomic and transcriptomic responses of these extremophytes to salt stress ^20–24^, molecular mechanisms determining how they have convergently achieved salt adapted growth remains unknown.

In this study, we used a multi-omics experimental design integrating transcriptomics, metabolomics, and ionomics to investigate the coordinated cellular responses and the associated genetic mechanisms employed by *S. parvula*, *E. salsugineum,* and *A. thaliana* to survive salt stress at variable success levels. We examined the spatio-temporal coordination of multiple genetic pathways used by the three models at different salt stress intensities. Both extremophytes showed distinct molecular phenotypes suggesting induction of complementary cellular processes that used core pathways present in all plants, but with modifications to those in ways that optimized balance between stress tolerance and growth.

## Results

Previous studies using either *S. parvula* or *E. salsugineum* in a comparative study with *A. thaliana* have successfully used 150 mM NaCl to elicit salt stress responses in the extremophyte while avoiding induction of immediate tissue necrosis in *A. thaliana* in the short term ^6^. We grew *S. parvula, E. salsugineum*, and *A. thaliana* hydroponically for 28 days; then transferred them to a medium supplemented with 150 mM NaCl for comparative studies including *A. thaliana* (Figure S1A). *Arabidopsis thaliana* did not show severe stress symptoms until four days of exposure to salt. Therefore, we decided to capture the short-term effects of salt at 0, 3, 24, and 72 hr after salt treatment to include timepoints that would precede the onset of stress phenotypes (Figure S1). Additionally, for the extremophytes, 250 mM NaCl was used to further examine their salt stress responses. We used the initial three timepoints to detect immediate responses of the ionome and the transcriptome to salt and used 0 hr and the latter two timepoints to detect the subsequent metabolome alterations (Figure S1B).

### High tissue tolerance to Na accumulation vs limiting Na accumulation within tissues

All three species accumulated Na as the duration of the treatment or the concentration of NaCl increased (Figure 1A). *Arabidopsis thaliana* shoots showed a 19-fold increase of Na compared to control within 3 hours of exposure to salt. It accumulated more Na in shoots than in roots and did not limit Na accumulation in roots as much as *S. parvula*. Tissue accumulation of Na was lowest in *S. parvula* compared to the other two species. Interestingly, *E. salsugineum* allowed higher Na accumulation in both roots and shoots compared to *S. parvula,* at levels similar to those observed for *A. thaliana* roots under salt stress (Figure 1A). This suggests that *S. parvula* limited total Na accumulation in tissues while *E. salsugineum* showed high tissue tolerance to Na accumulation, a trend maintained for 250 mM NaCl treatments in the two extremophytes (Figure 1A).

**Figure 1.**
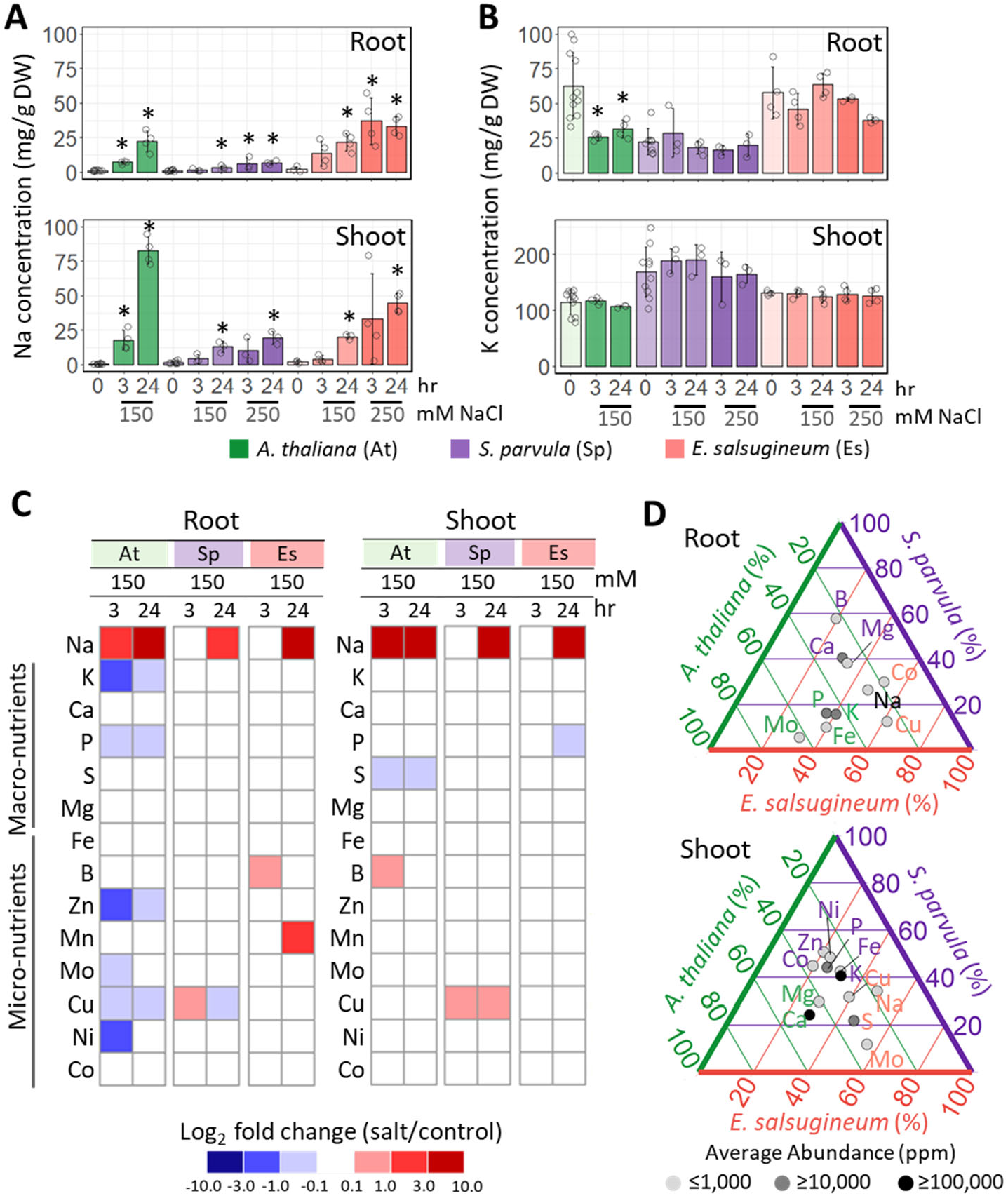
Sodium accumulation and its effect on nutrient balance in the extremophytes compared to *A. thaliana* under salt stress. [A] Na and [B] K content in roots and shoots. Data are mean ± SD (n ≥ 4, at least 4 plants per replicate). Significant differences between treatment groups were determined by ANOVA with Tukey’s test applied to within species comparisons. Asterisks indicate significant difference (*p* ≤ 0.05) between the treated samples and their respective controls. Open circles indicate biological replicates. [C] Salt stress-induced changes in macro-and micro-nutrient content. [D] Percent abundance of nutrients that showed significant differences in abundance at control conditions among the three species. Significant difference in elemental abundance was determined by one-way ANOVA with post-hoc Tukey’s test at *p* ≤ 0.05. The three axes of the ternary plots are marked with *A. thaliana, S. parvula*, and *E. salsugineum*. The gridlines in species-designated colors point to % abundance of the element from the corresponding species over the total abundance of that element from all three species. Thus, gridlines of each color nearest to each data point lead to the relative % abundance in each species.

High Na levels interfere with K and other nutrient uptake in plants (Munns and Tester, 2008). We examined whether Na accumulation triggered nutrient imbalance in any of the test species by quantifying the abundance of 13 plant nutrients during salt treatments. The K level significantly dropped within 3 hr in *A. thaliana* roots, but did not change in either extremophyte, regardless of the duration and intensity of the Na treatments (Figure 1B). In shoots, we did not observe any significant changes in K levels by salt treatments in any of the three species. When we examined the profiles of all quantified nutrients, shoots exhibited minimal changes in response to salt stress compared to roots (Figure 1C). Intriguingly, *A. thaliana* roots showed significant depletion of 6 out of 13 nutrients under salt treatments whereas both extremophytes showed minimal nutrient disturbances (Figure 1C and Table S1). We then examined nutrient compositions in the control samples (basal levels) to identify nutrients preferentially enriched in any of the species (Figure 1D and Table S1). Notably, Ca and Mg, which are known to regulate specificity of Non-Selective Cation Channels (NSCCs) ^25^, were found at higher basal levels in *S. parvula* roots compared to other species. Similarly, Cu and Co were high in *E. salsugineum* whereas Fe, P, K, and Mo were high in *A. thaliana*.

### Primary metabolite pools decreased from high basal levels in *E. salsugineum* but increased from low basal levels in *S. parvula* upon salt treatments

Salt stress requires metabolic adjustments in plants ^7^. Therefore, we examined the metabolite adjustment strategies of the two extremophytes in comparison to salt-sensitive *A. thaliana*. We quantified a total of 716 metabolites using GC-MS for each species (see Methods), of which 182 with known specific identities were referred to as “known metabolites’’ while the remaining were collectively referred to as “unknown metabolites’’ in this study (Table S2). The overall quantified metabolite pool in roots showed a relatively strong correlation in metabolite abundance between selected pairwise comparisons resulting in two distinct trends (Figure 2A). First, the control groups of *S. parvula* and *A. thaliana* exhibited the highest pairwise correlation, and this correlation decreased as the duration of salt treatment increased (Figure 2A, upper panel). Second, *S. parvula* and *E. salsugineum* profiles were moderately correlated at control conditions, but upon salt treatments, this correlation became stronger, indicating that the majority of metabolites in the extremophytes adjusted to similar levels in the roots. At 150 mM NaCl, this increase in correlation between *S. parvula* and *E. salsugineum* was only apparent at 72 hr, but at the higher NaCl concentration of 250 mM, their root metabolite profiles were strongly correlated even at 24 hr (Figure 2A, lower panel). In contrast to roots, the shoot metabolomes of *S. parvula* and *E. salsugineum* remained divergent following salt treatments (Figure S2A).

**Figure 2.**
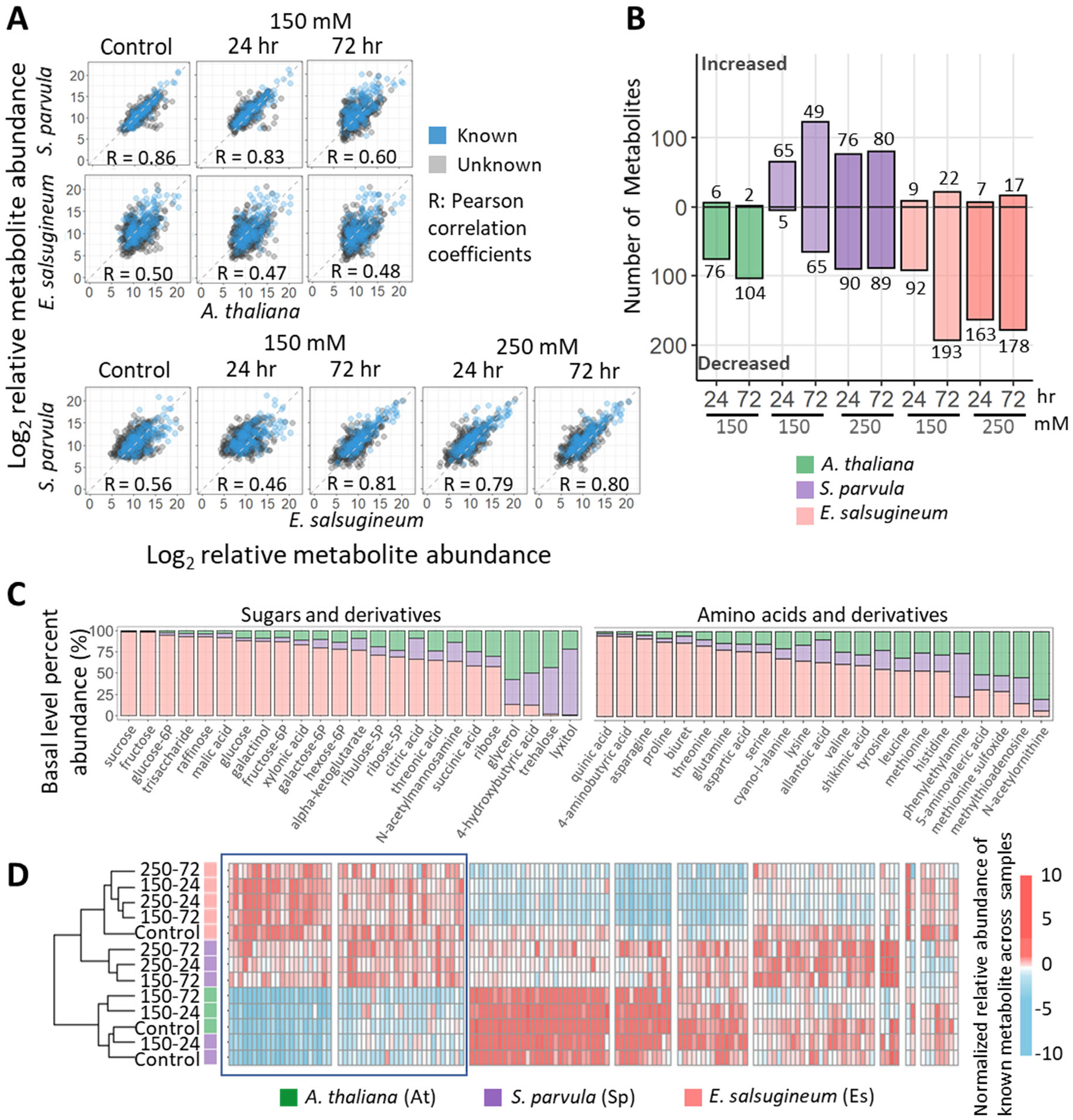
Overall metabolic preparedness and adjustments in the roots of the extremophytes compared to *A. thaliana*. [A] Pairwise Pearson correlations of the abundance of 716 quantified metabolites across all conditions and species. Blue dots indicate 182 metabolites with known structures, and grey dots represent 534 unknown metabolites. [B] Number of metabolites that significantly changed in abundance (DAMs) in each species compared to its respective controls. [C] Basal level percent abundances of sugars, amino acids, and their derivatives in the three species. [D] Hierarchical clustering of known metabolites shown in [B and C]. Blue box highlights clusters of interest. Columns indicate metabolites and rows indicate samples. Sample names for treated conditions are given as treatment concentration and duration, separated by a dash. Treatment concentrations are 150 and 250 mM NaCl; treatment durations are 24 and 72 hr. Significant differences were determined by one-way ANOVA with post-hoc Tukey’s test at *p* ≤ 0.05 (n = 4, at least 4 plants per replicate).

Under salt treatments, metabolite abundances in both *A. thaliana* and *E. salsugineum* roots largely decreased while *S. parvula* metabolite abundances increased (Figure 2B). This trend for *S. parvula* was not upheld in shoots, where metabolite abundances in all three species predominantly decreased (Figure S2B). We next examined if the dynamically changing metabolites were categorically associated with sugars and amino acids often known for their roles as organic osmolytes (Slama et al., 2015). Figure 2C shows percent abundance among the three species at control levels in roots, for all sugars, amino acids, and their immediate derivatives (see Methods) quantified in our study that showed a significant difference in abundance among the three species. Even before the salt treatment, *E. salsugineum* had accumulated much higher levels for most of these metabolites than the other species. For instance, the abundance of sucrose in *E. salsugineum* accounted for 98.6% of the combined sucrose abundance in the roots of all three species (Figure 2C) and this was 155-and 131-fold higher than that in *A. thaliana* and *S. parvula*, respectively (Table S2). Similarly, glucose, raffinose, fructose, and proline were more abundant in *E. salsugineum* compared to the other species (Figures 2C and Table S2).

We checked whether one extremophyte maintained a higher basal abundance for key metabolites associated with osmoregulation while the other extremophyte actively induced those when exposed to salt, eventually converging to a metabolic status distinct from that of *A. thaliana* under salt stress in roots, as we observed in Figure 2A. We clustered abundance profiles of all known metabolites exhibiting significant changes in abundance at the basal level among the three species or in at least one stress condition in any species (Figure 2D). Two readily identifiable clusters (box outlined in Figure 2D) showed lower metabolite abundances in *A. thaliana* and control/150 mM-24 hr *S. parvula* samples compared to *E. salsugineum*, but each of those metabolites increased exclusively in *S. parvula* upon prolonged/higher salt treatment to match the constitutive high level found in *E. salsugineum*. These clusters were enriched in metabolites known for their role as osmoprotectants or antioxidants ^26^, including sucrose, fructose, glucose, GABA, proline, and dehydroascorbic acid (full list in Table S3). These results indicate a basal level metabolic “preparedness” in *E. salsugineum* roots compared to active induction of many osmoprotectants and antioxidants in *S. parvula* when responding to salt treatments. On the other hand, *A. thaliana* lacks neither a preadapted-nor a dynamic-strategy found in the two extremophytes.

### *Arabidopsis thaliana* showed stronger transcriptomic responses compared to the extremophytes during salt stress

Previous studies have reported that nearly 20% of the *A. thaliana* transcriptome responds to salt stress ^21, 23^. We tested if the transcriptomic responses from *A. thaliana* aligned more with either extremophyte or if the extremophytes showed a largely overlapping response that was distinct from *A. thaliana*. The overall root and shoot transcriptome profiles were grouped into species-tissue clusters (Figure 3A). These dominant species-level distinctions in transcript profiles were further illustrated by pairwise correlations of entire transcriptome profiles (Figure S3). Unlike metabolite profiles (Figures 2A), none of the transcriptome comparisons showed strong correlations between species in any pairwise comparison (Figure S3).

**Figure 3.**
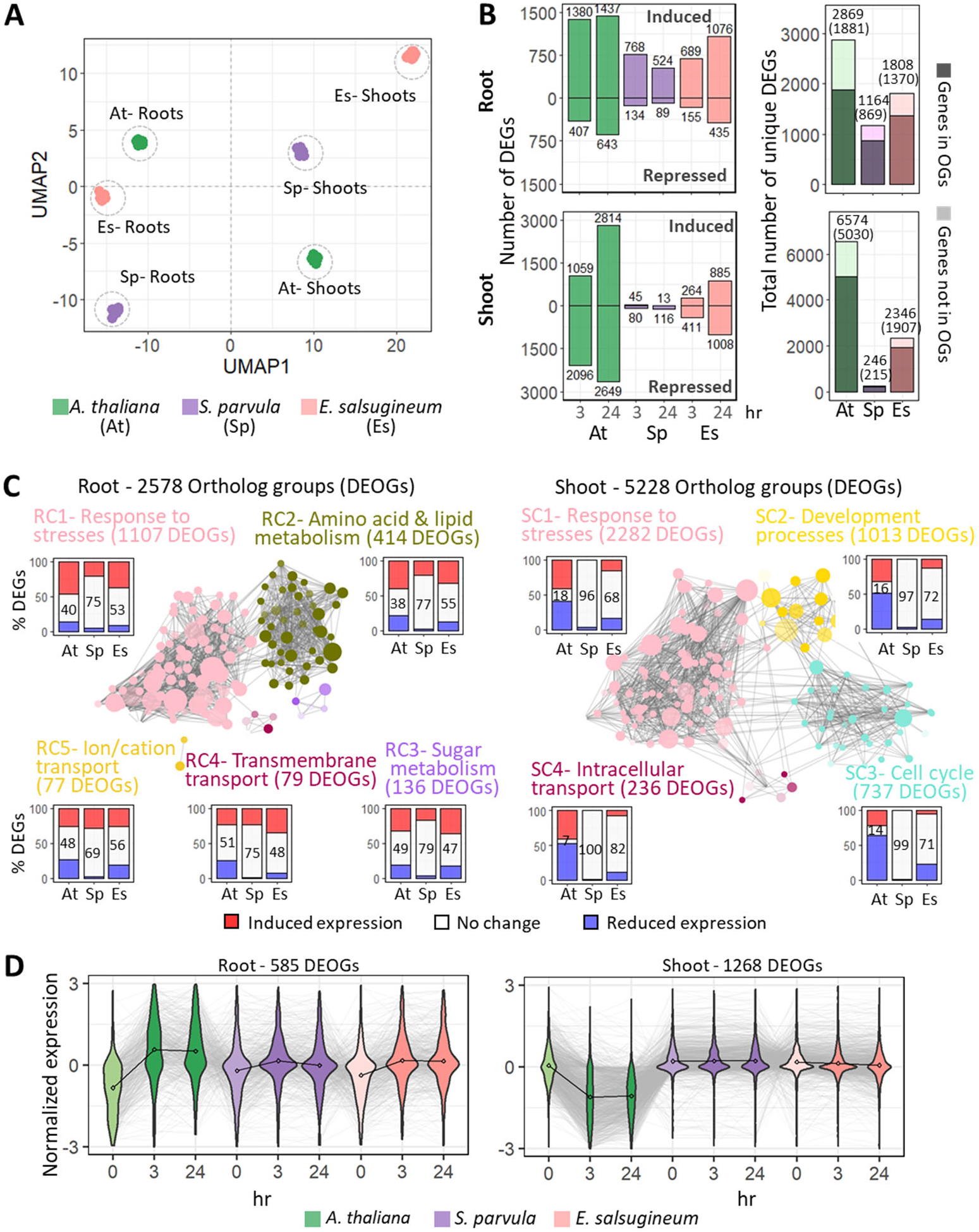
Transcriptomic overview of *Schrenkiella parvula, Eutrema salsugineum,* and *Arabidopsis thaliana* in response to salt. [A] Overall clustering of transcriptomes in shoots and roots from all replicates in all conditions using one-to-one ortholog groups (OGs). [B] Number of differentially expressed genes (DEGs) in response to salt treatments. DEGs were identified using DESeq2 at *p-*adj ≤ 0.01 (n = 3, at least 4 plants per replicate). [C] Functional clusters enriched among differentially expressed ortholog groups (DEOGs) that included a DEG from at least one species in roots (RC, root cluster) and shoots (SC-shoot cluster). Functional annotations were based on GO annotations assigned to DEOGs. The bar graph assigned to each cluster represents percent allocation of induced (red) and suppressed (blue) DEGs with the percentage of genes that did not significantly change in each species given in the white space. [D] Dominant gene expression clusters of DEOGs across time points and species from roots and shoots that included an *A. thaliana* ortholog that was either induced or suppressed in response to 150 mM NaCl treatment. The average gene expression at each condition is indicated by a dot and connected by a black line for each species. Individual gene expression is indicated by grey lines.

Among differentially expressed genes (DEGs) in the roots of all three species, we observed more transcripts induced by salt treatments than suppressed (Figure 3B, left panels). The majority of these DEGs were identified in ortholog groups (OGs) (see Methods) (Figure 3B, right panels, and Table S4). *Arabidopsis thaliana* showed the largest number of DEGs in response to salt stress in both roots and shoots. Interestingly, the *A. thaliana* shoot transcriptome had more than twice the number of DEGs observed in the root transcriptome in response to salt stress (Figure 3B), although the nutrient profiles under salt stress in shoots showed minimal changes compared to roots (Figure 1C). *Arabidopsis thaliana* also had thousands of salt-responsive DEGs compared to remarkably fewer DEGs in the extremophytes especially in shoots (Figure 3B).

We next examined if transcriptomic changes associated with cellular homeostasis in *A. thaliana* entailed processes already maintained or induced in the extremophytes during salt treatments. This search was done at two levels. First, we clustered all 1-to-1 orthologs based on their basal expression levels (from control samples); filtered the clusters to show expression patterns that differed at least in one species compared to the other two; and identified enriched functional processes in each cluster (Table 1). Responses associated with abiotic stress were among the most highly representative processes in each of these clusters. Second, we investigated the changes in transcriptomic profiles during salt treatments by identifying functional clusters enriched among all ortholog groups (OGs) that included at least one DEG in a species, considering roots and shoots separately (Figure 3C and Table S5 and S6). Response to stress formed the largest functional cluster including the highest number of OGs in both roots and shoots (RC1 and SC1 in Figure 3C), which were further sub-clustered to highlight various salt responsive pathways mediated by auxin and ABA regulation and oxidative stress responses (Figure S4). Amino acid metabolism (RC2) and sugar metabolism (RC3) formed the next two largest root clusters followed by transmembrane transport (RC4) and ion transport (RC5) (Figure 3C). In shoots, the second and third largest functional clusters (SC2 and SC3 in Figure 3C) suggested cellular processes involved in plant development and growth/cell cycle, in which the majority of the *A. thaliana* orthologs were salt-repressed. The extremophytes showed enrichment of similar functional processes associated with abiotic stresses when treated with the 250 mM NaCl concentration (Figure S5).

**Table 1.**
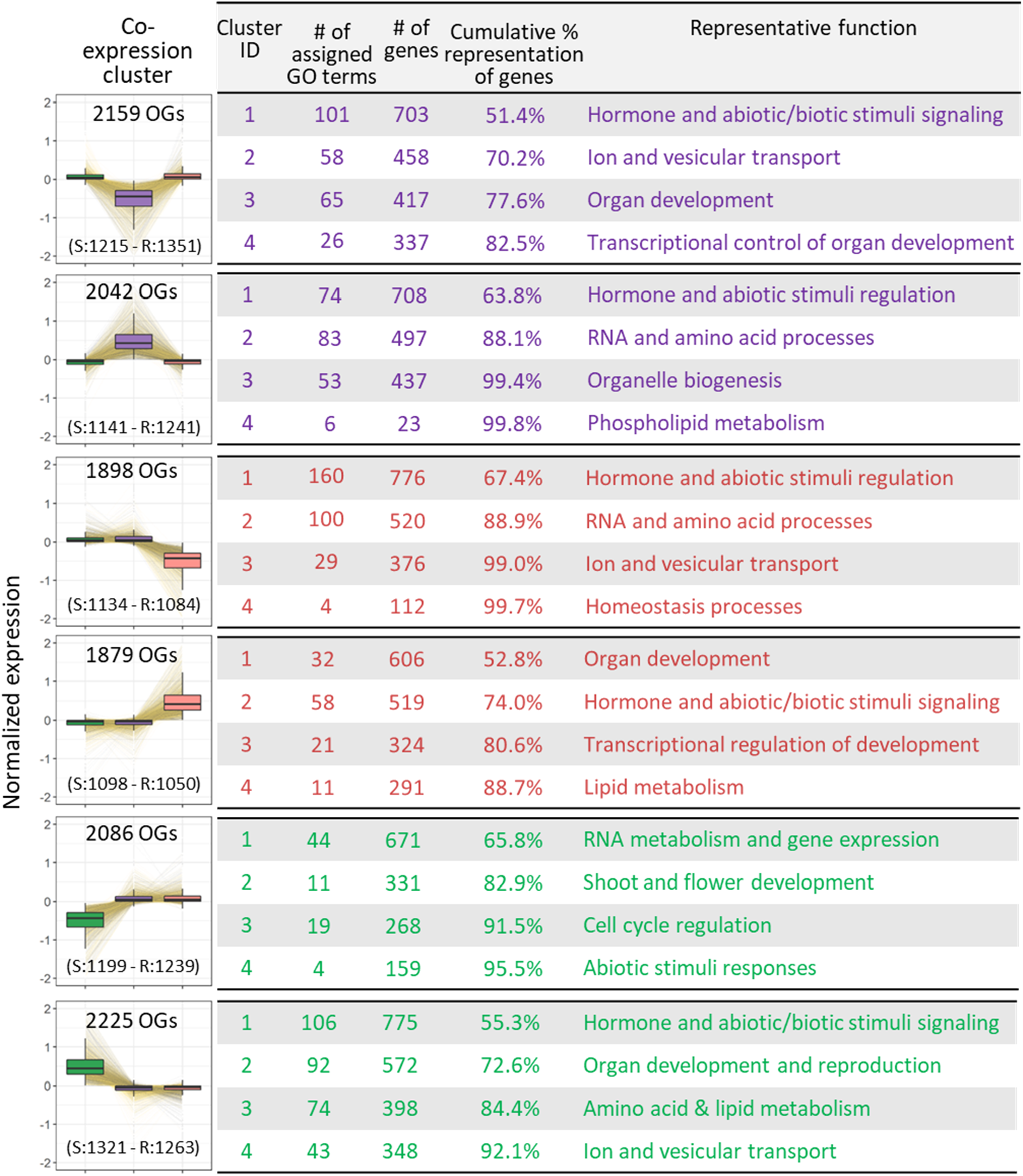
Cellular processes enriched among ortholog clusters identified based on their basal expression levels in shoots (S) and roots (R) of *S. parvula* (purple), *E. salsugineum* (red), and *A. thaliana* (green).

| Co-expression cluster | Cluster ID | # of assigned GO terms | # of genes | Cumulative % representation of genes | Representative function |
| --- | --- | --- | --- | --- | --- |
| 2159 OGs<br>(S:1215 - R:1351) | 1 | 101 | 703 | 51.4% | Hormone and abiotic/biotic stimuli signaling |
|  | 2 | 58 | 458 | 70.2% | Ion and vesicular transport |
|  | 3 | 65 | 417 | 77.6% | Organ development |
|  | 4 | 26 | 337 | 82.5% | Transcriptional control of organ development |
| 2042 OGs<br>(S:1141 - R:1241) | 1 | 74 | 708 | 63.8% | Hormone and abiotic stimuli regulation |
|  | 2 | 83 | 497 | 88.1% | RNA and amino acid processes |
|  | 3 | 53 | 437 | 99.4% | Organelle biogenesis |
|  | 4 | 6 | 23 | 99.8% | Phospholipid metabolism |
| 1898 OGs<br>(S:1134 - R:1084) | 1 | 160 | 776 | 67.4% | Hormone and abiotic stimuli regulation |
|  | 2 | 100 | 520 | 88.9% | RNA and amino acid processes |
|  | 3 | 29 | 376 | 99.0% | Ion and vesicular transport |
|  | 4 | 4 | 112 | 99.7% | Homeostasis processes |
| 1879 OGs<br>(S:1098 - R:1050) | 1 | 32 | 606 | 52.8% | Organ development |
|  | 2 | 58 | 519 | 74.0% | Hormone and abiotic/biotic stimuli signaling |
|  | 3 | 21 | 324 | 80.6% | Transcriptional regulation of development |
|  | 4 | 11 | 291 | 88.7% | Lipid metabolism |
| 2086 OGs<br>(S:1199 - R:1239) | 1 | 44 | 671 | 65.8% | RNA metabolism and gene expression |
|  | 2 | 11 | 331 | 82.9% | Shoot and flower development |
|  | 3 | 19 | 268 | 91.5% | Cell cycle regulation |
|  | 4 | 4 | 159 | 95.5% | Abiotic stimuli responses |
| 2225 OGs<br>(S:1321 - R:1263) | 1 | 106 | 775 | 55.3% | Hormone and abiotic/biotic stimuli signaling |
|  | 2 | 92 | 572 | 72.6% | Organ development and reproduction |
|  | 3 | 74 | 398 | 84.4% | Amino acid & lipid metabolism |
|  | 4 | 43 | 348 | 92.1% | Ion and vesicular transport |

We have identified ortholog clusters showing salt-responsive co-expression across time points within tissues among species. We presented the two most dominant co-expression clusters in roots and shoots (Figure 3D and Table S7). In roots, the largest co-expression cluster showed that *A. thaliana* orthologs were substantially salt-induced, compared to a much smaller magnitude of induction seen in the extremophyte orthologs (Figure 3D, left panel). In shoots, the largest co-expression cluster included *A. thaliana* orthologs suppressed in response to salt, while expression of the extremophyte orthologs was stable (Figure 3D, right panel).

### The predominant transcriptomic signature in roots supports divergent auxin-dependent root growth during salt stress

In all three species, response to stress was the most significantly enriched function in roots among ortholog sets showing either different basal-level expression (Table 1) or salt-responses (Figure 3C) among species. The largest representative function among all subclusters were response to hormones in all ortholog sets, especially in roots (Table 1, Figures S4A, S5A, and S5C). Hormonal regulation plays an important role in salt stress responses ^27^. Furthermore, our comparative ortholog expression profiling suggested that regulation of hormone signal transduction differed among the target species (Figures 4A, S6, and Table S8). Genes mediating auxin response via auxin/indole acetic acid (Aux/IAA) repressors ^28^ and the type 2C protein phosphatases (PP2Cs) that negatively regulate abscisic acid (ABA) signaling ^29^ showed contrasting salt-responsive expression among the three species. Specifically, in response to salt, 15 out of 24 *AUX/IAAs* and *PP2Cs* orthologs in *A. thaliana* and *E. salsugineum* were significantly induced whereas none of those orthologs in *S. parvula* showed any change (Figures 4A, S6, and Table S8). When the basal expression of these orthologs were examined, 11 out of 17 *E. salsugineum AUX/IAA* orthologs and 5 out of 7 *S. parvula PP2C* orthologs showed higher basal expression compared to their respective orthologs in the other species (Figure 4A bottom panel, and Table S8). In contrast, expression patterns of orthologs encoding regulators of other steps in the auxin and ABA signaling pathway as well as other hormone signaling pathways showed much fewer or no differences among the three species (Figures S6, and Table S8).

**Figure 4.**
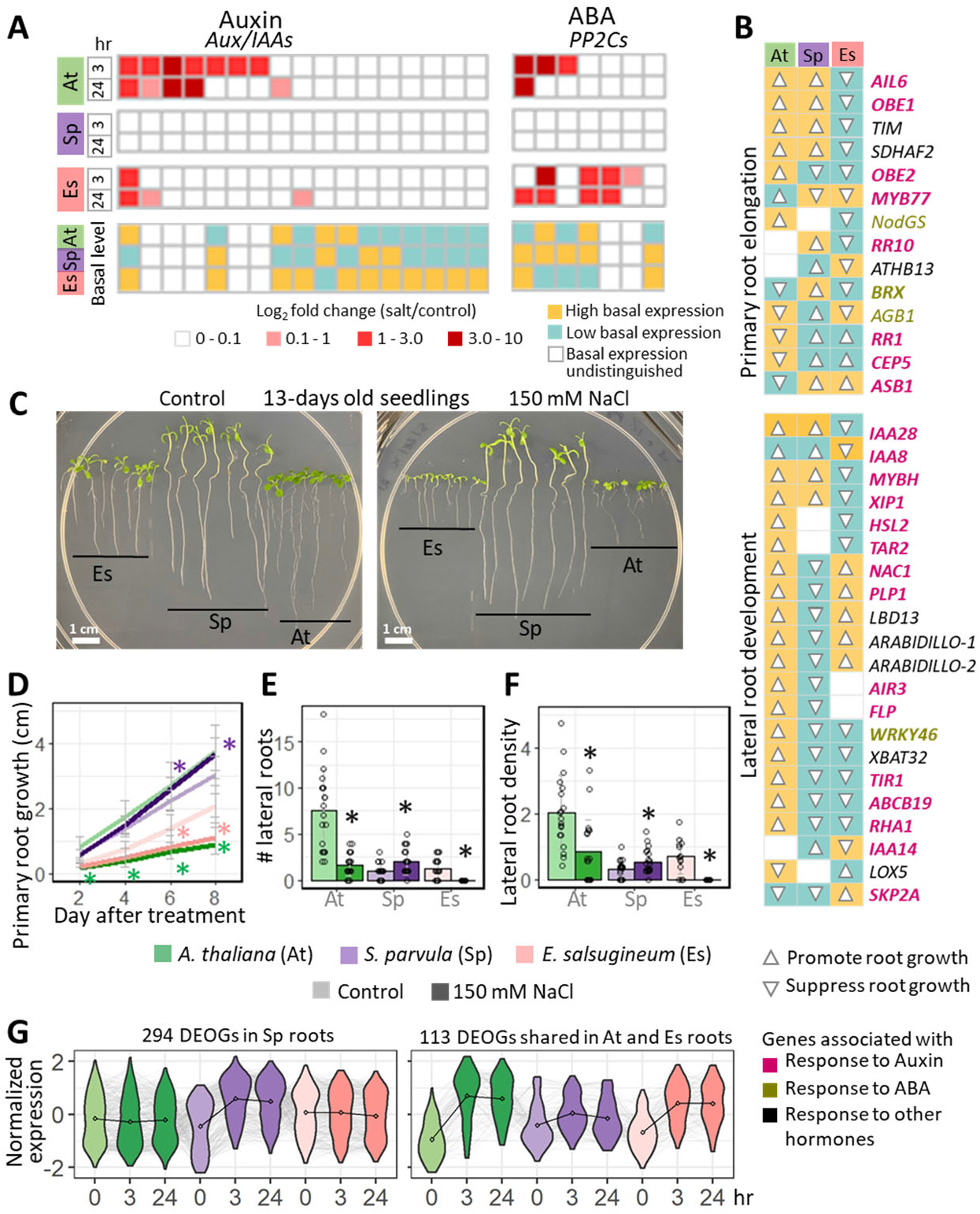
Root growth responses in line with gene expression associated with auxin and ABA signaling in *Arabidopsis thaliana, Schrenkiella parvula,* and *Eutrema salsugineum* when treated with salt. [A] Salt-induced changes in expression of auxin/indole acetic acid (Aux/IAA) repressors and type 2C protein phosphatases (PP2Cs) that regulate auxin and ABA signaling. [B] Curated genes from lateral root development (GO:0048527) and primary root development (GO:0080022) used to infer root growth phenotypes based on functional genetic studies in *A. thaliana*. Ortholog expression levels are categorized as high (yellow), low (turquoise), and indistinguishable (no-color) based on their relative basal expression among the three species. Their inferred effect on root growth is indicated by the arrowheads (up for promoting growth and down for suppressing growth). Genes associated with auxin, ABA, and other hormones were labeled in pink, gold, and black respectively. [C] Root growth examined with and without salt. [D] Primary root growth, [E] number of lateral roots, and [F] lateral root density of 13-day-old seedlings that were treated with 150 mM NaCl for 8 days. Data are given as mean ± SD (n = 3, at least 7 plants per replicate). Open circles indicate individual measurements. Asterisks indicate significant differences (*p* ≤ 0.05) between treated samples and their respective control samples, determined by one-way ANOVA with post-hoc Tukey’s test. [G] Gene co-expression clusters of differentially expressed ortholog groups (DEOGs) that showed similar expression pattern in *A. thaliana* and *E. salsugineum* roots compared to a different pattern observed for *S. parvula* under salt stress. Left panel includes DEOGs that are uniquely upregulated in *S. parvula*; right panel includes DEOGs that are co-upregulated in *A. thaliana* and *E. salsugineum*. The average gene expression at each condition is indicated by a dot and connected by a black line for each species. Individual gene expression is indicated by grey lines.

Auxin and ABA suppress root elongation and lateral root initiation during salt stress in *A. thaliana* ^30, 31^. Therefore, we wanted to investigate if auxin and ABA signaling pathways that showed significant alterations in their expression profiles in the target species could lead to distinct root phenotypes indicated by genetic studies in *A. thaliana.* We first selected all known genes involved in primary root development (GO:0080022) and lateral root development (GO:0048527) and checked for functionally verified phenotypes associated with those genes that describe root growth in *A. thaliana*. We curated a list of orthologs that directly led to root growth promotion or suppression if they are highly expressed or suppressed (see full list of references used for the selected genotype-phenotype associations in Table S9). Next, we assigned a binary category of promotion or suppression of root growth to each ortholog based on their high or low basal expression levels in one species compared to the other two. The final set of 35 single copy orthologs were assigned to a root growth map (Figure 4B) and a gene network for lateral root development (Figure S7, modified from De Rybel et al., 2010^32^; Banda et al., 2019^33^). We found 27 orthologs in *S. parvula* that suggested increased primary root elongation or suppressed lateral root initiation, while 23 orthologs in *E. salsugineum* suggested slower primary and lateral root growth when compared to the remaining two species (Figures 4B and S7). Contrastingly, *A. thaliana* showed 25 orthologs that supported its fast primary root growth and increased lateral root number. Notably, the majority (71%) of these 35 orthologs were annotated as auxin-responsive genes (Figure 4B).

In line with the observed differences in expression of orthologs associated with root development (Figure 4B and S7), *S. parvula* and *A. thaliana* seedlings indeed have comparable primary root lengths at control conditions, longer than primary roots in *E. salsugineum* (Figures 4C and D). Moreover, *S. parvula* showed uncompromised primary root growth compared to *E. salsugineum* and *A. thaliana* when treated with salt for an extended time (Figures 4C, D, and S8). *Schrenkiella parvula* and *Eutrema salsugineum* seedlings also had fewer lateral roots compared to *A. thaliana* at control conditions (Figure 4E). However, lateral root growth, assessed using total lateral root number and density during a week-long salt treatment, indicated that *S. parvula* not only sustained uninterrupted root growth but also induced lateral root initiation albeit lower initial numbers during salt treatments, in contrast to the responses observed for *A. thaliana* and *E. salsugineum* (Figures 4E, F, and S8).

To further identify orthologs in *S. parvula* that support its unique pattern of uninterrupted primary root growth and lateral root initiation during salt stress, we checked co-expression clusters that included differentially expressed ortholog groups (DEOGs) in roots. We found two clusters where *S. parvula* showed a distinct expression pattern compared to the other two species (Figure 4G). The first cluster showed salt-induced expression unique to *S. parvula.* This accounted for 11% (294 ortholog groups) of all DEOGs in roots (Table S10). The second cluster with 113 OGs (5% of all root DEOGs) showed induction of genes in response to salt stress in *A. thaliana* and *E. salsugineum* while the orthologs in *S. parvula* did not alter their expression (Figure 4G). Interestingly, within these two clusters, nearly 50% of orthologs did not have a GO annotation describing a specific pathway or molecular function implying the substantial unexplored functional gene space of these orthologs (Table S10).

### Transcriptome-metabolome coordination in extremophytes provides protection from salt stress

There were 7,852 differentially expressed ortholog groups (DEOGs) and 634 differentially abundant metabolites (DAMs) collectively in the three species in response to all salt treatments (Figures 2 and 3, Table S2 and 4). The difference in % change in response to salt between DAMs and DEGs was the lowest in *A. thaliana*, while in extremophytes metabolic responses were relatively larger compared to the overall transcriptomic response during salt treatments (Figures 5A and S9A). This indicates that there is more preparedness at the transcriptome level to respond to salt along with a dynamically responsive metabolome in the extremophytes than in *A. thaliana*.

**Figure 5.**
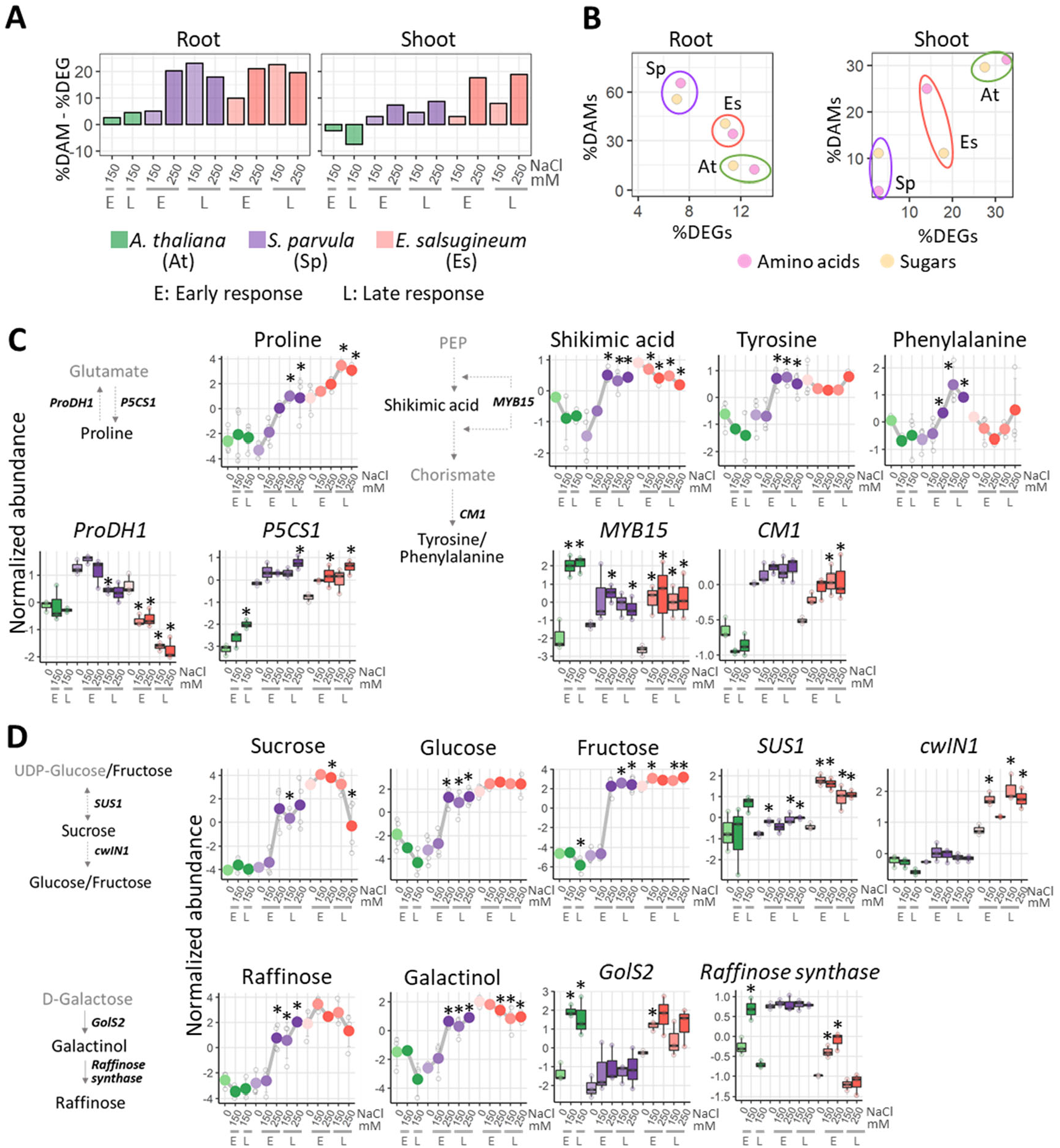
Coordination between differentially expressed genes (DEGs) and differently abundant metabolites (DAMs) during responses to salt in *Arabidopsis thaliana, Schrenkiella parvula,* and *Eutrema salsugineum*. [A] The difference in the percentage of DEGs and DAMs in response to salt treatments. [B] Percentage of DEGs and DAMs involved in the metabolism of amino acids, sugars, and their immediate derivatives. DEGs were categorized based on their GO annotations: GO:0006520 for cellular amino acid metabolism and GO:0005975 for carbohydrate metabolism. Selected pathways in [C] amino acid and [D] sugar metabolism with concordant DAMs and DEGs. Line graphs represent normalized log2 relative metabolite abundance. Boxplots represent normalized gene expression values. Center line in the boxplots indicates median; box indicates interquartile range (IQR); whiskers show 1.5 × IQR. Asterisks indicate significant difference between the treated samples and their respective controls (n = 3-4). Early (E) and late (L) responses for transcripts refer to 3 and 24 hr, respectively. E and L responses for metabolites refer to 24 and 72 hr, respectively. Metabolites are shown in the backbone of the pathway while genes encoding for key enzymes/transcription factors are placed next to the arrows. Metabolites that were quantified in the current study are given in black while those not quantified are in grey.

The second and third largest functional clusters over-represented among differentially expressed ortholog groups (DEOGs) in roots were amino acid and sugar metabolism (550 DEOGs) (Figure 3C, RC2 and RC3). Additionally, the overall metabolite response indicated a high correlation of the metabolomes between the two extremophytes *S. parvula* and *E. salsugineum* during salt treatments, and the DAMs shared between the extremophytes were enriched in amino acids and sugars (Figure 2). Collectively, these findings led us to examine whether a coordinated transcriptomic response (i.e. DEGs associated with amino acid and sugar metabolism) support a salt-induced metabolic response (i.e. DAMs that are amino acids and sugars) in the extremophytes. In line with the global trend (Figure 5A), we observed fewer DEGs associated with amino acid and sugar metabolism in both extremophyte roots than in *A. thaliana* roots, but more amino acid and sugar metabolites were found as DAMs in extremophytes in roots (Figure 5B).

Given the established role of amino acids and sugars as organic osmoprotectants during salt stress ^26^, we focused on highly responsive DAMs that are also osmoprotectants and the key genes that encode proteins involved in the synthesis of those metabolites (Figures 5C and D, and S9A). Proline was one of the most significant DAMs found in the roots of both extremophytes. It was not only found at a higher basal level in the extremophytes than in *A. thaliana*, but also its abundance further increased with salt treatments specifically in the extremophytes (Figure 5C). Concordant with this increase in proline abundance, both extremophytes increased their transcript levels for *pyrroline-5-carboxylate synthetase* (*P5CS1*), which encodes the primary proline biosynthesis enzyme, and decreased expression for *Pro-dehydrogenase* (*ProDH1*), which encodes an enzyme involved in proline catabolism (Figures 5C and S9B). Additionally, shikimic acid-derived amino acids, tyrosine, and phenylalanine (precursors of phenylpropanoids) showed higher basal level abundances or significant inductions in roots of both extremophytes compared to *A. thaliana* (Figure 5C). Interestingly, genes coding for the enzymes directly involved in the conversion of phosphoenolpyruvate to shikimic acid such as DAHPS1, DAHPS2, DHQS, DHQ, SK1, ESPS, and CS were not detected as DEGs in any of the three species (Figure S9C). However, *MYB15,* a master regulator of shikimic acid biosynthesis pathway (Chen et al., 2006), was highly induced in all three species (Figure 5C). *Chorismate mutases 1* (*CM1*), coding for the enzyme involved in the first committed step of tyrosine and phenylalanine biosynthesis was further induced in *E. salsugineum*, while it was constitutively expressed at a high level in *S. parvula* (Figure 5C).

*Eutrema salsugineum* maintained a higher basal abundance than the other two species for multiple sugars in roots that are commonly used as organic osmolytes in plants, while these metabolites increased their abundances in response to salt in *S. parvula* (Figure 5D). The genes encoding enzymes involved in the biosynthesis of these metabolites such as *sucrose synthase 1* (*SUS1*), *cell wall invertase 1* (*cwIN1*), *galactinol synthase 2* (*GolS2*), and *raffinose synthase* were mostly induced under salt or constitutively highly expressed in the extremophytes (Figure 5D). A similar concordant alignment of DAMs to DEGs, or maintenance of high constitutive abundance of metabolites with high basal expression of genes were also found for metabolites and their associated genes in the extremophytes, known for their functions as antioxidants (for example, dehydroascorbic acid pathway highlighted in Figure S9D).

### Coordination between nutrient balance and gene expression associated with ion transport

Ion and membrane transport represented the two remaining major functionally enriched clusters in roots that included 156 DEOGs (Figure 3C, RC4 and RC5). As noted earlier, nutrient balance in the extremophytes was maintained during salt treatments, unlike that in *A. thaliana* (Figure 1C). To examine how the transcriptomic response specifically supported better nutrient balance in the extremophytes compared to *A. thaliana* (Figure 1), we first investigated all 148 transporter genes associated with Na/K transport (based on Araport11 (Cheng et al., 2017)) in the following four categories: (1) aquaporins (*PIPs*, *NIPs*, *TIPs*, and *SIPs*), (2) cation transporters (*CAXs*, *NHXs*, *KEAs*, *CHXs*, *KUPs*, *HKT1*, and *TRH1*), (3) non-selective cation channels (*CNGCs* and *GLRs*), and (4) K channels (*SKOR, GORK, AKTs* and *KCOs*). We found 109 corresponding one-to-one ortholog groups (OGs), among which 64 OGs were differentially expressed at basal expression among the three species (Table S11). We focused on the OGs encoding transporters and channels that were differentially expressed at basal levels in both extremophytes compared to *A. thaliana* (Figure 6A). Genes encoding transporters known to exclude Na from the cell (*SOS1*), transporters involved in Ca uptake and transport (*CNGC1*, *CNGC12*, *CAX1*, and *CAX5*), K channel (*KAT1*), and endosomal K transporter (*KEA5*) that aid in pH and ion homeostasis ^34, 35^ showed higher basal expression in the two extremophytes than in *A. thaliana*, while genes involved in K efflux/Na influx to the cell or water transport such as *TIP1;1*, *PIP2;2*, *SIP1;2, SIP2;1*, *KUP5/6/10/11*, and *NHX3* were lower expressed in the extremophytes (Figure 6A, Table S11). *NHX1/2*, the main transporters involved in Na sequestration into the vacuole ^36^, showed higher basal level expression in *E. salsugineum* compared to the other two species (Table S11). This observation aligns with *E. salsugineum* showing a higher Na sequestration (also higher tissue tolerance) than *S. parvula* and *A. thaliana* (Figure 1A).

**Figure 6.**
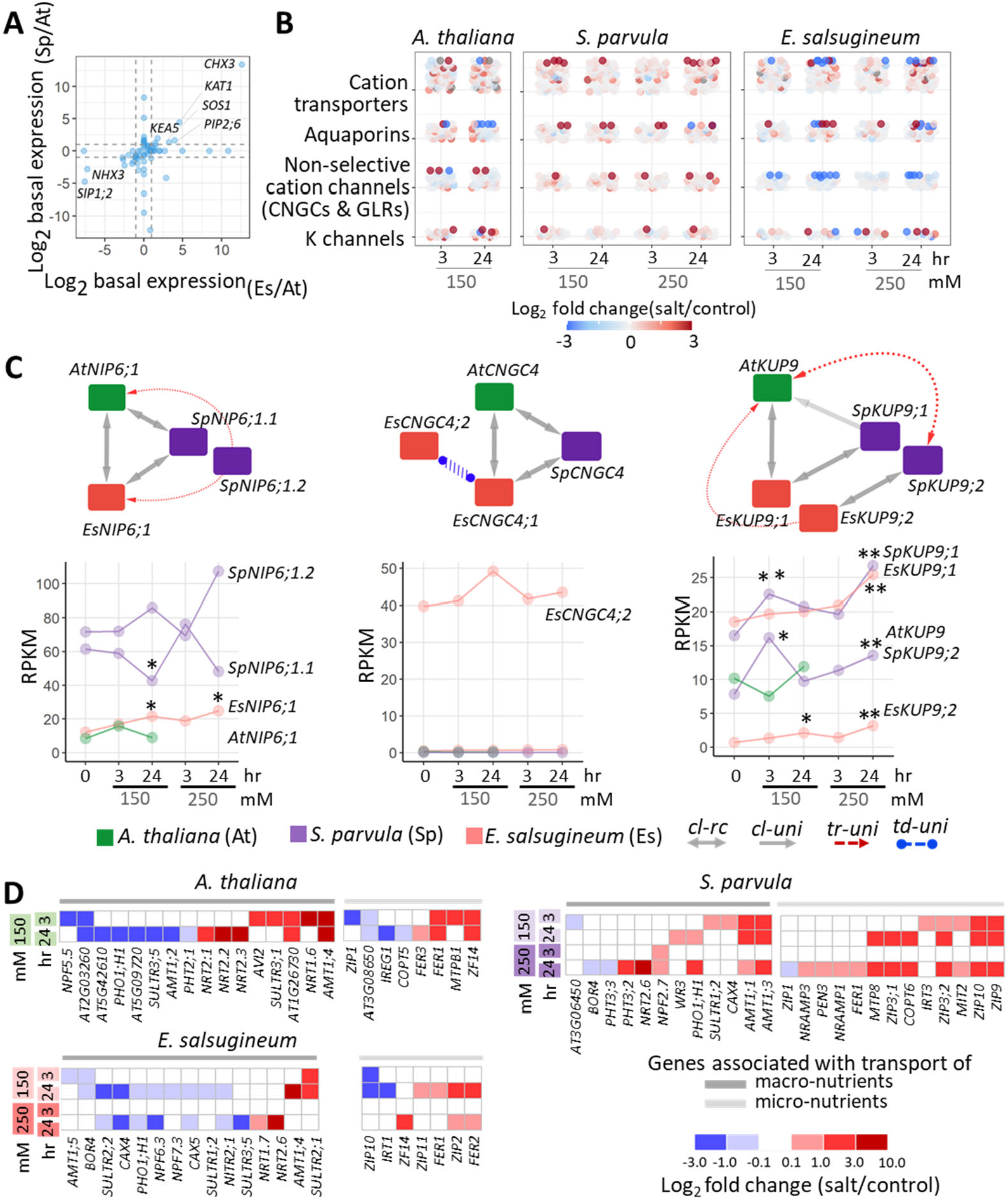
Transcriptomic support for Na and nutrient transport in *Arabidopsis thaliana, Schrenkiella parvula,* and *Eutrema salsugineum*. [A] Basal expression comparison of genes encoding putative Na/K transporters and channels in the roots. The dashed lines indicate 2-fold differences in gene expression between the extremophytes and *A. thaliana* to highlight genes either highly suppressed or highly induced in the extremophytes in roots at control conditions. [B] Expression changes of transporters/channels associated with Na/K transport activity during salt treatments. [C] Selected cation transporters that showed a higher gene copy number in extremophytes and their expression during salt treatments. Nodes assigned with species colors show the ortholog network. Edges show homology relationships, either co-linear reciprocal orthologs (cl-rc), co-linear unidirectional orthologs (cl-uni), transposed-unidirectional duplicates (tr-uni), or tandem duplicated-unidirectional paralogs (td-uni). RPKM: reads per kilobase of transcript per million reads mapped. Asterisks indicate significant difference between the salt treated samples and their respective controls (n = 3, at least 4 plants per replicate) at \**p*-adj ≤ 0.05 and \*\**p*-adj ≤ 0.01. [D] Salt-induced changes in the expression of genes associated with the transport of nutrients altered in the ionome (identified in Figure 3C). Sample names are shown on the left indicating treatment concentrations (150 and 250 mM NaCl) and durations (3 and 24 hr).

We then examined how these genes in different transporter classes changed in expression in the three species when treated with salt (Figure 6B and Table S12). Interestingly, the majority of the genes across the four categories were either significantly induced or constant in expression during salt treatments in *S. parvula*, whereas *A. thaliana* and *E. salsugineum* had a mixed response (Figure 6B). *Eutrema salsugineum* mostly repressed the expression of genes encoding non-selective cation channels (e.g. *GLRs*) in response to all salt treatments (Figure 6B and Table S12). Other observable trends included the induction of *CHX17* (functions in K uptake) identified as the only induced transporter in all three species in our selected set; a 50-fold induction of *CHX2* (involved in Na transport to vacuole) in A*. thaliana* (Table S12); and *GORK* as the only K channel up-regulated in *E. salsugineum* under salt treatment while maintaining high basal level expression in *S. parvula*.

We next expanded the focus on the transporter genes to include copy number variation among species and expression partitioned to paralogs present in each species (Figure 6C). We searched for transporter gene orthologs with increased copy numbers in the extremophytes and found, *SpNIP6;1*/*2* in *S. parvula*, *EsCNGC4;1*/*2* in *E. salsugineum*, *SpKUP9;1*/*2* and *EsKUP9;1*/*2* in both extremophytes (Figure 6C, top panels, and Table S13). All extremophyte paralogs either showed differential expression in response to salt or high constitutive expression compared to *A. thaliana* (Figure 6C, bottom panels).

Finally, we examined the expression profiles of all non-Na/K transporters involved in nutrient uptake that were included in DEOGs in Figure 3C functional clusters RC4 and RC5 (Figure 6D and Table S13). Notably, *S. parvula* induced orthologs encoding a large and diverse set of transporters involved in nutrient uptake upon salt treatments, while *A. thaliana* and *E. salsugineum* showed salt-induction of a limited group of genes associated with Fe, Zn, and Cu uptake (e.g. *FER1/*2/*3*, *MTPB1*, *ZF14*, *ZIP2/11*) (Figure 6D).

## Discussion

While the direct flow of Na ions into plant tissue is unavoidable in saline soils, plants show adaptations at varying degrees to limit accumulation of Na or mitigate cellular toxicity caused by Na in tissues ^2, 3, 7^. Figure 7 summarizes the most notable ionomic, metabolomic, transcriptomic, and phenotypic adjustments that distinguish the three model plants used in this study when responding to salt treatments. Overall, *S. parvula* and *E. salsugineum* show similar levels of salt tolerance, but they achieve this tolerance by different means. *Schrenkiella parvula* exhibits uncompromised primary root growth and nutrient uptake while limiting Na uptake. This coincides with an increase in amino acids and sugars that can serve as nitrogen-carbon sources, organic osmolytes, and antioxidants beneficial during salt stress. In contrast, *E. salsugineum* does not restrict Na accumulation in roots, and grows much more slowly under salt stress. It accumulates sugars and amino acids at remarkably high levels compared to the other two species. We find that the two extremophytes regulate different ortholog groups in the same pathways in response to salt stress in multiple comparisons. The divergent responses observed across multiple –omics data lead to a convergently salt tolerant phenotype in the extremophytes (Figure 7).

**Figure 7.**
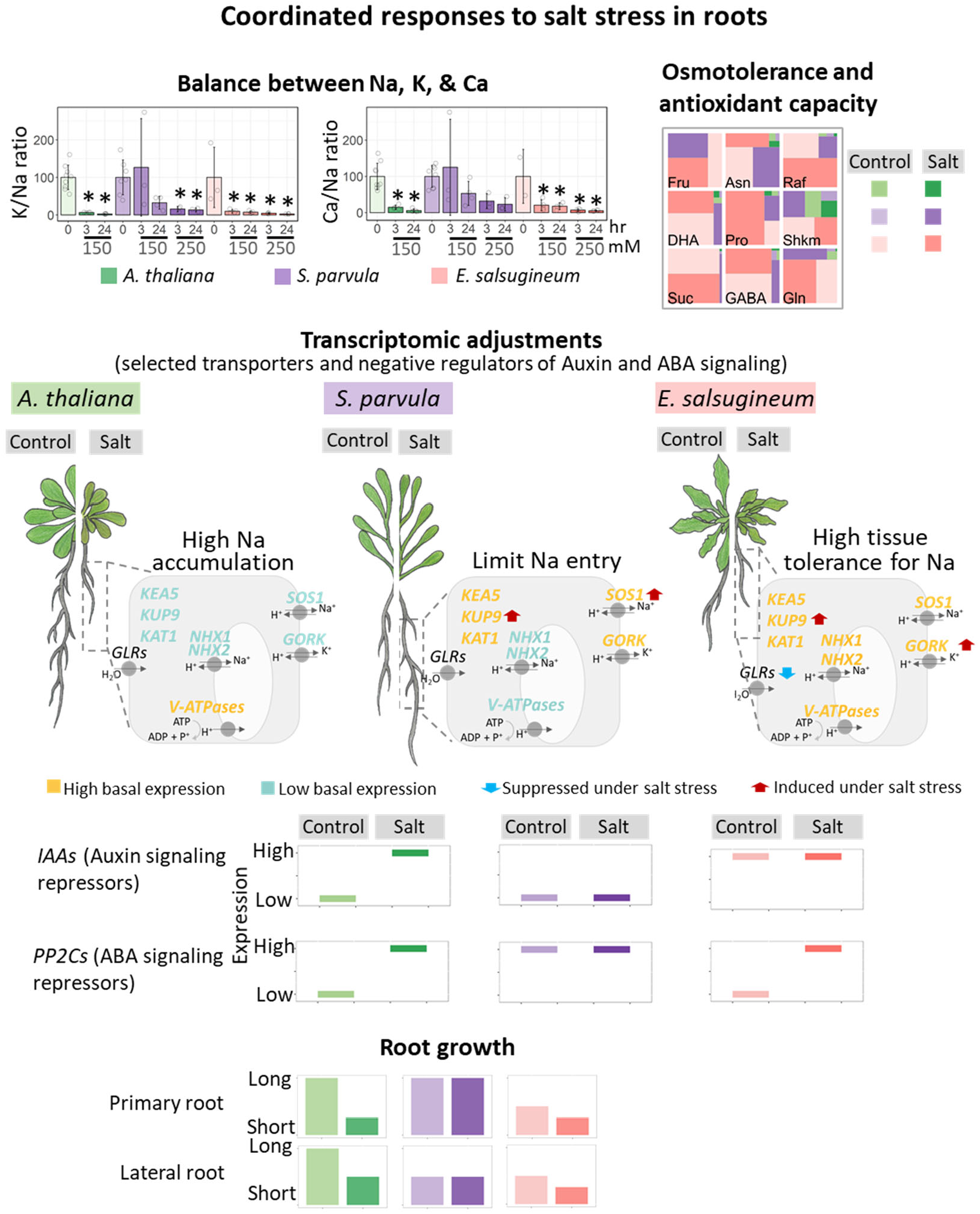
Focal responses to salt stress prevalent in roots of extremophytes compared to *A. thaliana* through ionomic, metabolomic and transcriptomic adjustments coupled to root growth. Ionomic balance is highlighted for Na, K, and Ca. Data are mean ± SD (n ≥ 4, at least 4 plants per replicate). Significant differences between treatment groups were determined by ANOVA with Tukey’s test within species. Asterisks indicate significant difference (*p* ≤ 0.05) between the treated samples and their respective controls. Open circles indicate the biological replicates. Top 9 metabolites enriched in the extremophytes. Fru, fructose; Asn, asparagine; Raf, raffinose; DHA, dehydroascorbic acid; Pro, proline; Shkm, shikimic acid; Suc, sucrose; GABA, gamma-aminobutyric acid; Gln, glutamine. Relative expression changes associated with ion transport and stress signaling. *KEA5*, K efflux antiporter 5; *KUP9*, K uptake permease 9; *KAT1*, K channel in *Arabidopsis thaliana* 1; *GLRs*, glutamate-like receptors; *NHX1*/*2*, Na/H exchanger1/2; *V-ATPases*, vacuoles-ATPases; *SOS1*, salt overly sensitive 1; *GORK*, gated outwardly-rectifying K channel. Relative root growth compared to control root growth in *A. thaliana*.

### Divergent response to Na accumulation and convergent outcome in maintaining nutrient balance during salt stress in extremophytes

Unlike *A. thaliana*, which fails to maintain its nutrient balance under salt stress as the Na content within tissues increases (Figure 1A-C), the two extremophytes present two distinct paths for regulating Na levels in roots. *Schrenkiella parvula* regulates Na accumulation to maintain it at levels observed for control conditions, while preventing the loss or maintaining the uptake of K and Ca. In contrast, *E. salsugineum* does not show such a restriction to Na accumulation in roots but reaches the same outcome as *S. parvula*, such that a significant loss of K and Ca is prevented.

Regulation of Ca and K is a critical requirement for ion homeostasis and response to excess Na ^3, 7^. Ca activates Na efflux via the SOS2-SOS3 signaling pathway, which in turn activates the plasma membrane localized Na-exporter, *SOS1* ^37^. Additionally, Ca regulates the plasma membrane localized non-selective cation channels (NSCCs), which are reportedly a main entry point of Na into roots ^38, 39^. Genes encoding NSCCs were highly suppressed in *E. salsugineum* during salt stress, consistent with previous findings ^40^ (Figure 6B and 7). However, counterintuitively, these *NSCCs* were induced or unaltered in *S. parvula* during salt treatments (Figure 6B), despite it having a relatively smaller Na content than salt-treated *E. salsugineum* (Figure 1A). The high basal level of Ca in *S. parvula* roots may alleviate Na-induced cellular toxicity by increasing the selectivity of NSCCs to limit Na entry ^39^ (Figure 1D).

Halophytes are known for their ability to maintain high K/Na ratios when exposed to salt, although the species-dependent ratios may vary among halophytes ^3, 41, 42^. Under salt stress, Na-induced membrane depolarization at the root epidermis could be reversed by the activation of the K outward rectifying channel, *GORK,* by releasing cytosolic K as seen in *A. thaliana* root hairs ^43, 44^. *GORK* expression in roots was induced in *E. salsugineum* when exposed to 250 mM NaCl and maintained at a high basal level in *S. parvula,* compared to *A. thaliana* (Tables S4 and 12). Therefore, GORK channels may help to prevent the membrane from further depolarization in extremophytes at high salinities. It is unclear how the extremophytes regulate their K transporters to allow K uptake and simultaneously prevent excessive leakage of K during salt stress. The extremophytes seems to have an alternative co-regulation of Na and K transporters different from *A. thaliana*. One such modified pathway in the extremophytes points to a possible synergistic activity between GORK and KAT1 (K channels) synchronized with Na exclusion mediated by SOS1 to regulate cytosolic K levels, pH, and ion homeostasis when exposed to salt. Coordinated expression of *GORK* and *KAT1* has been reported for guard cells in *A. thaliana* ^45, 46^. Compared to *A. thaliana*, *KAT1* is highly expressed at basal expression in the roots of both extremophytes (Figure 6A). Notably, the expression of these K channels is regulated by ABA and auxin signaling pathways, which are modified in both extremophytes compared to *A. thaliana* in conjunction with root growth modulation in response to salt (Figures 4, S7, and 7) ^47^. Intracellular K transport, as seen with *KEA5* (a K transporter in the trans-Golgi network) and *KUP9* (mediates K and auxin transport from the endoplasmic reticulum) ^34, 48^ appear to be differently regulated between the extremophytes and *A. thaliana* (Figures 6A and 7). It would be interesting to investigate whether the *KUP9* duplication in the extremophytes has led to divergent salt-responsive regulation in auxin homeostasis, and consequently lead to resilient root growth.

### Metabolic preadaptation or a dynamic response to achieve salt-adaptation in extremophytes

Sugars, amino acids, or their derivatives are used in all plants for osmoregulation during salt stress ^26^. The most striking metabolic feature among the three species in our study is the extraordinarily high levels of sugars and amino acids in *E. salsugineum* even in control conditions compared to the other two species (Figures 2C and 7). The *E. salsugineum* metabolite profiles we observed are consistent with earlier studies that examined selected metabolites in *E. salsugineum* ^49–51^. *Eutrema salsugineum* leaves and seedlings are metabolically preadapted to salt stress ^20–22^. Our results reinforce this view and extend these observations to roots (Figures 2C and 7). In contrast, *S. parvula* maintains much lower levels of sugars and amino acids comparable to levels seen in *A. thaliana* in control conditions and increases their levels in response to salt to the high basal level present in *E. salsugineum* (Figures 2 and 7). Complementary to this dynamic metabolic response, *S. parvula* exhibits a higher degree of transcriptome preadaptation to salt stress, having fewer DEGs in both roots and shoots in comparison to the other two species (Figures 3B and 5A).

Proline is among the most studied osmoprotectants and antioxidants in plants ^3, 52^. While the key proline biosynthesis gene, *P5CS1*, was induced by salt in all three species, *ProDH1*, encoding an enzyme that degrades proline, was suppressed only in the extremophytes, coincident to significant proline accumulation during salt stress (Figure 5C). *ProDH1* is suppressed by sucrose in *A. thaliana* ^53^. Our results suggest a novel regulatory mode for proline accumulation via high sucrose levels (facilitated by induced *SUS1* which is regulated by osmotic stress independent of ABA ^54^ that may lead to the suppression of *ProDH1*, and thereby allowing proline accumulation in the extremophytes (Figure 5D).

High levels of sugars and amino acids may serve as a carbon or nitrogen source to maintain growth when photosynthesis and nitrogen acquisition decrease with abiotic stress ^27, 51, 55^. *Eutrema salsugineum* roots grow much slower than *A. thaliana* and *S. parvula* primary roots under control conditions (Figure S8). A high proportional allocation of sugars such as sucrose and raffinose together with amino acids known for nitrogen storage, especially proline, is a common trait associated with slow-growing plants ^56, 57^. Some of these metabolites that are high at basal levels may have added benefits for serving as osmoprotectants or antioxidants when a slow-growing annual such as *E. salsugineum* experiences high salinity levels. *Schrenkiella parvula* dynamically accumulates these metabolites during salt treatments, without an indication of compromised growth, to match the levels maintained in *E. salsugineum* (Figures 4 and 5) ^58^.

### Divergent root transcriptional networks with convergent stress-resilient growth in extremophytes

*Schrenkiella parvula* and *Eutrema salsugineum* have lower lateral root densities compared to *A. thaliana* (Figure 4F). Auxin is a major hormone that regulates root architecture including lateral root development ^59^. Lateral roots are initiated by binding of auxin to its receptor TIR1, resulting in degradation of the auxin signaling repressors, Aux/IAAs ^60^. In both extremophyte, the *TIR1* orthologs are suppressed in control conditions coincident to suppression of multiple *Aux/IAAs* and other downstream transcription factors in control or salt-treated conditions, a pattern collectively supporting the observed limited lateral root initiation ^33^ (Figures 4B and S7). The two extremophytes either show a slower root growth rate (in *E. salsugineum*) or higher primary root growth rate (in *S. parvula*) coupled with lower lateral root density, which may reduce the total contact area with salt. Interestingly, the transcriptional network mediated by auxin and other hormones in the extremophytes indicates multiple regulatory points for root growth modulation leading to different root architectures (Figure S7).

Molecular phenotypes suggesting transcriptional preadaptation to multiple abiotic stresses have been reported for *S. parvula* and *E. salsugineum* separately ^20, 21, 23, 61–63^. Our work indicates that orthologs of stress-associated functional gene clusters that exhibit transcriptional preadaptation in one extremophyte are in many instances dynamically induced by salt in the other extremophyte and ultimately match the expression level found in the preadapted species (e.g. Figure 4G). This highlights independent and divergent regulation of orthologs even between closely related extremophytes in response to the same salt treatments, demonstrating different adaptive strategies.

Our study highlights different combinatorial expression modules for stress-optimized growth used by these two extremophytes to adapt to high salinities. For example, the selective release of repression in auxin and ABA signaling pathways likely leads to different *S. parvula* and *E. salsugineum* root growth strategies absent in *A. thaliana* and needs further investigation to identify additional regulatory dependencies using targeted studies in the extremophyte models. Similarly, comparisons from multiple extremophytes can serve as training data to identify compatible regulatory pathways that can coexist but are currently absent in crops. Such pathways need to be evaluated for their functionality to metabolic cost to decide whether constitutive expression (as pre-adaptive traits) or induced expression (as dynamic responses) offer an optimum strategy depending on the stress being constant or intermittent in certain environments. The scarcity of halophytes implies a high metabolic cost and complex regulation required for salt adaptation enabling coordinated growth from cellular to whole plant level ^64^. Extremophytes provide a direct resource when modeling stress resilient growth using core pathways critical to deliver growth and survival during environmental stress ^65^. For sustainable global food security, our crops need to be diversified; less dependent on fresh water and high nutrient soils; while being adapted to varying levels of marginal soils with different salinities ^66, 67^. Therefore, basic research examining genetic regulation underlying stress-optimized growth will be a prerequisite when selecting new crops in light of a climate crisis.

## Materials and methods

### Plant growth and treatments

*Schrenkiella parvula* (ecotype Lake Tuz, Turkey; Arabidopsis Biological Resource Center/ABRC germplasm CS22663)*, Eutrema salsugineum* (ecotype Shandong, China; ABRC germplasm CS22504), and *Arabidopsis thaliana* (ecotype Col-0) seeds were surface-sterilized and stratified at 4 °C for 7 days (for *A. thaliana* and *S. parvula*) or 14 days (for *E. salsugineum*). Stratified seeds were germinated and grown in a hydroponic system as described by Conn et al., (2013)^68^ for transcriptomic, ionomic, and metabolomic experiments (Figure S1B). The plants were grown in aerated 1/5x Hoagland’s solution in a growth cabinet set to 22-23 °C, photosynthetic photon flux density at 80-120 mM m^-2^s^-1^, and a 12 hr light / 12 hr dark cycle. Fresh Hoagland’s solution was replaced every two weeks. Four-week-old plants were randomly placed in 1/5x Hoagland’s solution with and without NaCl for indicated duration in each experiment.

### Root growth analysis

Stratified seeds were geminated on 1/4x MS medium. Five-day old seedlings were transferred to 1/4x MS plates supplied with 150 mM NaCl. Root growth was recorded every two days for one week. Control and treated seedlings were scanned and analyzed using ImageJ ^69^ to quantify primary root length and number of lateral roots. Three biological replicates were used with 7 seedlings per replicate from each species.

### Elemental analysis

Elemental quantification was conducted for Na, K, Ca, P, S, Mg, Fe, B, Zn, Mn, Mo, Cu, Ni, and Co using Inductively Coupled Plasma - Mass Spectrometry (ICP-MS, Elan 6000 DRC-PerkinElmer SCIEX) at the US Department of Agricultural Research Service at Donald Danforth Plant Science Center. We used dried root and shoot samples from 3-4 biological replicates from each condition harvested at 3 and 24 hr (Figure S1) processed as described in Baxter et al., (2014)^70^. Changes in element contents were calculated as the log_2_ fold change of the element level in treated samples over that in control samples and visualized with the pheatmap in R. Significant differences between treatments within species were determined by one-way ANOVA followed by Tukey post-hoc test using agricolae in R with an adjusted *p*-value cutoff of 0.05. Basal level of each element per species was calculated as a % contribution from each species that added to 100% for each element, using the following formula: X_i_/ (At_i_+Sp_i_+Es_i_)*100 where *X_i_* represents the abundance of element *i* in *A. thaliana*, *S. parvula* and *E. salsugineum*, respectively. Only elements that showed significant differences in abundance among the three species were visualized on ternary diagrams using ggtern in R.

### Metabolite analysis

Untargeted high throughput metabolite profiling was conducted using gas chromatography-mass spectrometry (GC-MS) service at the West Coast Metabolomics Center, University of California Davis. Root and shoot samples in 4 biological replicates were harvested at 24 and 72 hr (Figure S1B), flash frozen in liquid N_2_, and processed as described in Fiehn, (2016)^71^ and quantified as described in Pantha et al., (2021)^63^.

Pearson correlation coefficients for each test condition were calculated between species using Log_2_ median-normalized relative metabolite abundances. Significant differences in metabolite abundances across treatments within species (differently abundant metabolites, DAMs) was determined by one-way ANOVA followed by Tukey post-hoc test using agricolae in R with an adjusted *p*-value cutoff of 0.05. Structurally annotated (known) metabolites were further categorized into functional groups according to the refmet database (https://www.metabolomicsworkbench.org/databases/refmet/index.php) and Kyoto Encyclopedia of Genes and Genomes (KEGG). Metabolites were clustered into amino acids, sugars, nucleic acids, and other organic acids. Derivatives or precursors of those were included in the same metabolite category if those metabolites were found to be within three steps of the main metabolite category identified in a KEGG pathway (Table S2). Basal level of each metabolite per species was calculated as a % contribution from each species that added to 100% for each metabolite as described for the elemental basal level calculation. Metabolites that showed significant differences in abundance at basal level or under salt treatments in at least one species were used for K-mean clustering (k= 9) ^72^. The metabolite profiles across samples were done using pheatmap in R.

### Orthologous group identification

Ortholog groups were identified using *S. parvula* gene models version 2.2 from Phytozome (https://phytozome-next.jgi.doe.gov/); *A. thaliana* gene models version 10 (TAIR10) from TAIR (https://www.arabidopsis.org/download/), and *E. salsugineum* gene models version 210.3 from CoGe (https://genomevolution.org/coge/). Reciprocal gene pairs between species were identified using blastp with default parameters and an e-value cutoff of 1E-5. These pairs were filtered using a custom python script to convert a blast tabular output into a table of one query-subject pair per line by consolidating High-scoring Segment Pairs (HSPs); and based on calculated proportion of query and subject covered by HSPs (coverage) and approximate proportion of identical sequences within HSPs (identity). Pairs with an alignment coverage smaller than 50% of the query or the subject were removed. This ortholog-pair list included pairs with the highest bit score optimized for highest % coverage, and % identity. Ortholog pairs were additionally filtered to exclude any orthologs that had a more than ± 30% length difference in the coding sequence between the two sequences. A total of 16,591 one-to-one ortholog groups were identified among the three species and used for all downstream analyses (Table S3).

### Gene expression profiling and analysis

Total RNA was extracted from control and treated samples at 3 and 24 hr with three biological replicates (Figure S1) using Qiagen RNeasy Plant Mini kit (Qiagen, Hilden, Germany), with an on-column DNase treatment. mRNA enriched samples were converted to libraries using True-Seq stranded RNAseq Sample Prep kit (Illumina, San Diego, CA, USA), multiplexed, and sequenced on a HiSeq4000 (Illumina) platform at the Roy K. Carver Biotechnology Center, University of Illinois at Urbana-Champaign. A minimum of >15 million 50-nucleotide single-end reads per sample were generated for a total of 78 RNAseq samples.

After quality checks, the reads were mapped to reference transcript model sequences of gene models in each species (described earlier) using Bowtie ^73^ with -m 1 --best -n 1 -l 50. A custom python script was used to count uniquely mapped reads for each gene model. Differentially expressed genes (DEGs) across treatments within each species were identified using DESeq2 ^74^ with an adjusted *p*-value ≤ 0.01. Genes were further filtered using the following criteria: 1) |log_2_ fold change| ≥ 1 or 2) |log_2_ fold change| ≥ 0.5 if normalized mean expression across samples ≥ 100. Reads per kilobase per million reads (RPKM) were calculated for each gene from the raw read counts for all samples. These RPKM values were log_2_-transformed and median-normalized when used in PCA-UMAP ^75^ and for co-expression cluster determinations. Co-expression clusters were identified using fuzzy K-mean clustering with a membership cutoff ≥ 0.4 or binary clustering based on log_2_ fold change. Log_2_ fold changes were computed for all ortholog groups (OGs) from three species, subsequently used to calculate Pearson correlation coefficients when comparing transcriptome profiles between species/samples. Ortholog groups where at least one of the genes in the group was identified as a DEG in any species under any stress condition was considered to be a differentially expressed ortholog group (DEOG).

To compare the basal expression levels of orthologs among species, we generated a raw count matrix based on reads uniquely mapped to coding sequences (CDS) for the 16,591 orthologs from control samples. Orthologs that are differently expressed between any two of the three species were identified using DESeq2 at an adjusted *p*-value ≤ 0.001 in a pairwise manner. We next assigned the expression of each gene within any given OG as High (H), Medium (M), and Low (L) based on their relative expression level, which resulted in 12 clusters. The six largest clusters in both roots and shoots highlighted the expression difference in one species compared to the other two species. OGs assigned to these clusters were represented by HLH, LHL, HHL, LLH, LHH, and HLL (expression profiles). Each expression profile was then subjected to functional gene enrichment analysis.

BiNGO ^76^ was used to identify Gene Ontology (GO) terms enriched in selected DEGs and DEOGs. To reduce the redundancy between enriched GO terms, we further grouped them into GO clusters using GOMCL ^77^ with settings for -Ct 0.5 -I 1.5 -Sig 0.05 -hm -nw -hgt -d -gosize 3500 -gotype BP. Sub-clustering of selected clusters was performed using GOMCL-sub with the same parameters.

OrthNets for selected genes were generated using the CLfinder-OrthNet pipeline with the reciprocal blastp results as input using default OrthNet settings ^78^, and visualized in Cytoscape.

### Curated gene sets for primary metabolism, hormone signaling, root development, and transporter functions

OGs and DEOGs that showed significant differences at basal level/control condition or under treatments were mapped to the KEGG pathways associated with primary metabolism, hormone regulation, or root development. Log_2_ fold changes were computed for all selected OGs and visualized with pheatmap in R. Genes involved in root development were identified from Gene Ontology annotations, GO:0048527 for lateral root development and GO:0080022 for primary root development. Genes coding for transport functions associated with K/Na transport were mined from Araport 11. To assess the correlation between changes at the transcriptomic level and the metabolic level, we calculated the percentage of DEGs and differently abundant metabolites (DAMs) in each species at early (3 hr for transcriptome and 24 hr for metabolome) and late (24 hr for transcriptome and 72 hr for metabolome) response to salt. Percentage of DEGs was calculated by dividing the number of DEGs by the total number of expressed genes (RPKM ≥ 1) in each species. Similarly, the percentage of DAMs was calculated by dividing the number of DAMs by the total number of quantified metabolites within a species. We extracted DEGs and DAMs related to amino acid and sugar metabolism based on GO (GO:0006520 for amino acid metabolism and GO:0005975 for carbohydrate metabolism). The count of DEGs and DAMs were separately normalized to the total number of genes and metabolites in each of the two categories.

## Data availability

All Illumina sequence data are deposited at National Center for Biotechnology Information BioProject PRJNA63667. Mapped RNAseq data can be browsed using genome browsers created for *Schrenkiella parvula* and *Eutrema salsugineum* at www.lsugenomics.org.

## Acknowledgement

This work was supported by the US National Science Foundation IOS-EDGE 1923589/20196, NSF-MCB-1616827, US Department of Energy BER-DE-SC0020358, Next-Generation BioGreen21 Program of Republic of Korea (PJ01317301) awards. Graduate students K.T., G.W. were supported by an Economic Development Assistantship from LSU. We thank the LSU High Performance Computing facility for providing computational resources; undergraduate students Jordan Vivien, Christine Tran, Ashley Doan, and Jason Garcia at LSU for assisting with plant growth and phenotyping; Drs. Alvaro Hernandez and John Cheeseman at University of Illinois for providing assistance with Illumina sequencing.

## Author contribution

K.T. prepared plant samples and conducted data analyses; K.T., G.W., D-H.O., J.L., A.S., and M.D. contributed to data interpretation. K.T. and M.D. wrote the manuscript with input from all co-authors who revised and approved the final manuscript. M.D. conceptualized and supervised the overall project.

## Supplementary figure captions

**Figure S1.**
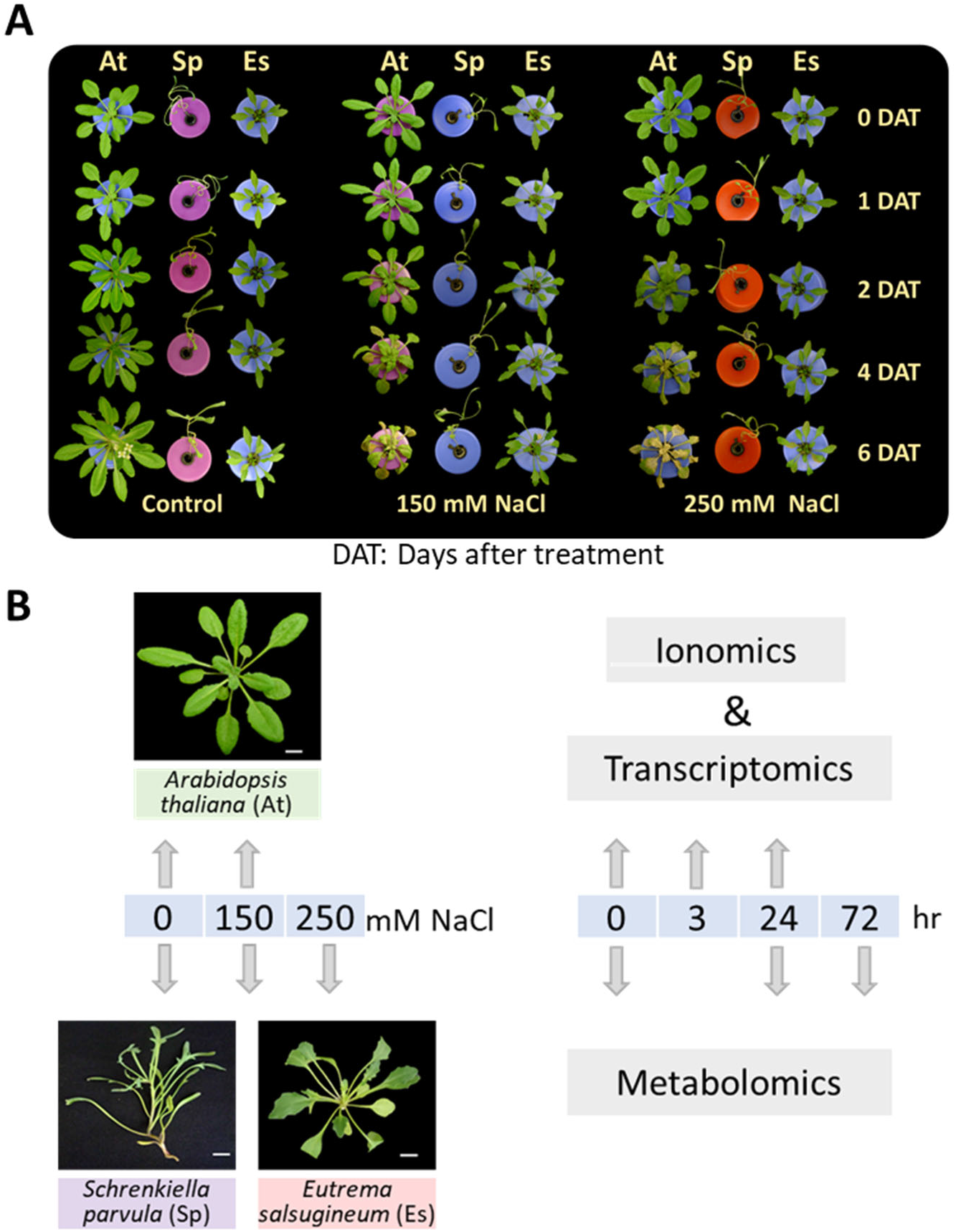
Effect of salt stress on *Schrenkiella parvula* (Sp), *Eutrema salsugineum* (Es), and *Arabidopsis thaliana* (At). [A] Four-week-old hydroponically grown plants and the salt concentrations examined in this study. DAT: Days after treatment. [B] Sampling scheme for ionomic, metabolomic, and transcriptomic profiling. There were at least 4 replicates per condition used for ionomic and metabolomic profiling, and 3 replicates per condition for transcriptomic profiling with at least 4 plants included per replicate.

**Figure S2.**
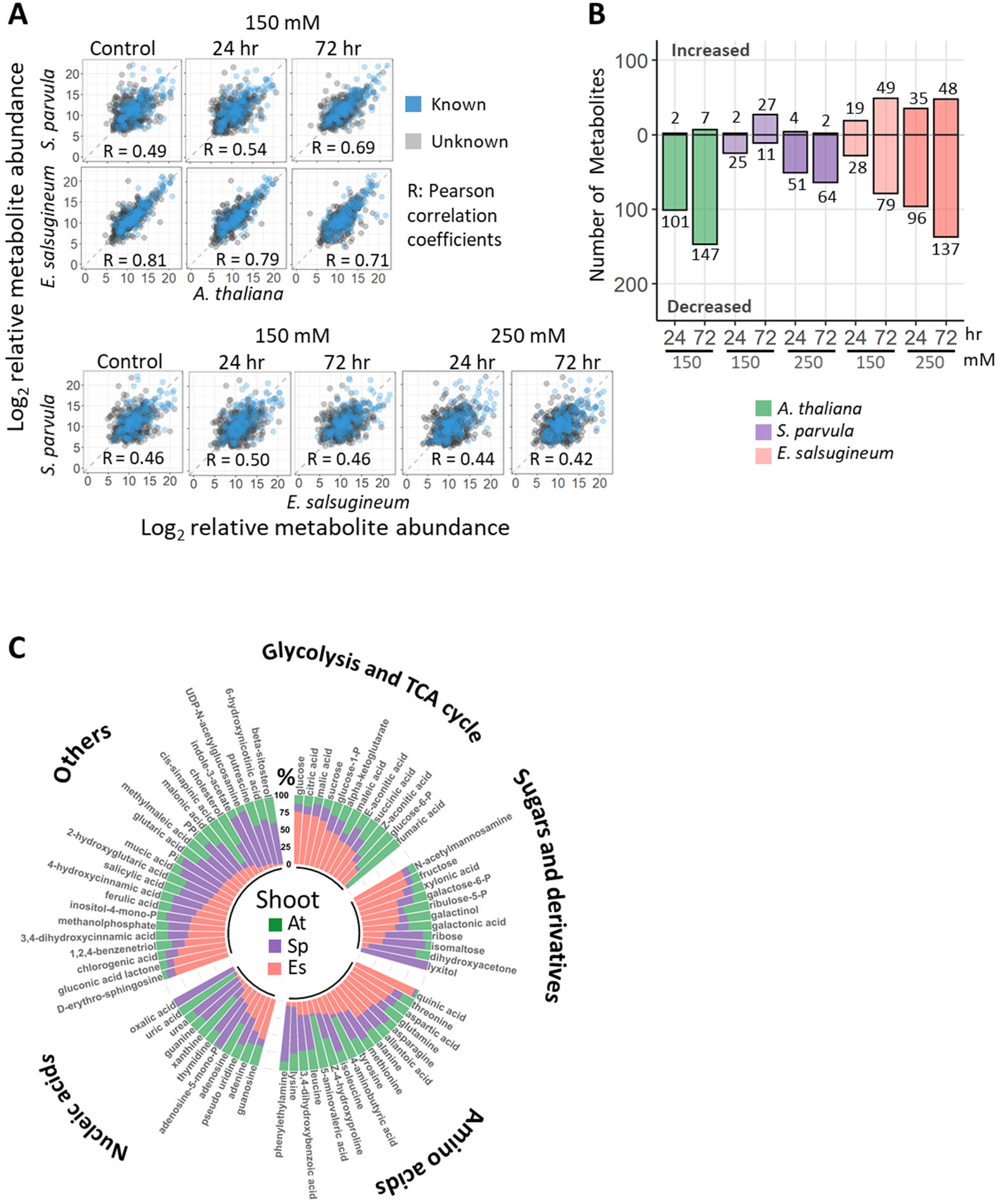
Overall metabolic adjustments in the shoots of the extremophytes compared to *A. thaliana*. [A] Pairwise Pearson correlations of the abundance of 716 quantified metabolites across all conditions and species. Blue dots indicate 182 metabolites with known structures, and grey dots represent 534 unknown metabolites. [B] Number of metabolites that significantly changed in abundance (DAMs) in each species compared to its respective control. [C] Basal level percent abundances of sugars, amino acids, and their derivatives in the three species. Significant differences were determined by one-way ANOVA with post-hoc Tukey’s test at *p* ≤ 0.05 (n = 4, at least 4 plants per replicate).

**Figure S3.**
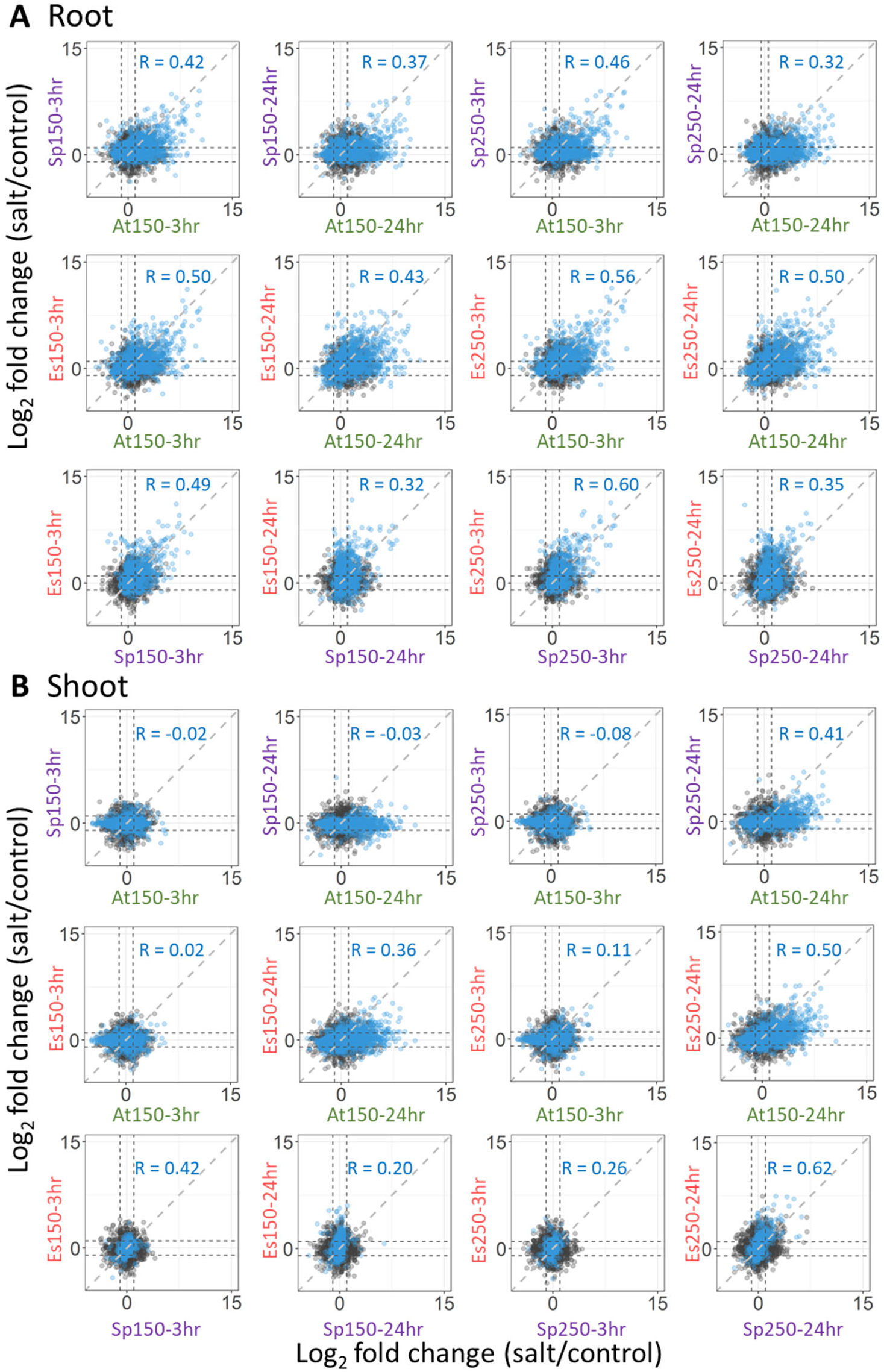
Pairwise comparison of transcriptomes of *Schrenkiella parvula*, *Eutrema salsugineum,* and *Arabidopsis thaliana* during response to salt treatments. Pearson correlations of [A] root and [B] shoot transcriptomes were calculated using differentially expressed ortholog pairs which included DEGs from at least one condition in one species. The axis labels are composed of species, treatment concentrations, and treatment durations. Axis color codes represent *S. parvula* (Sp) in purple, *E. salsugineum* (Es) in red, and *A. thaliana* (At) in green. Treatment concentrations include 150 and 250 mM NaCl; treatment durations include 3 and 24 hr.

**Figure S4.**
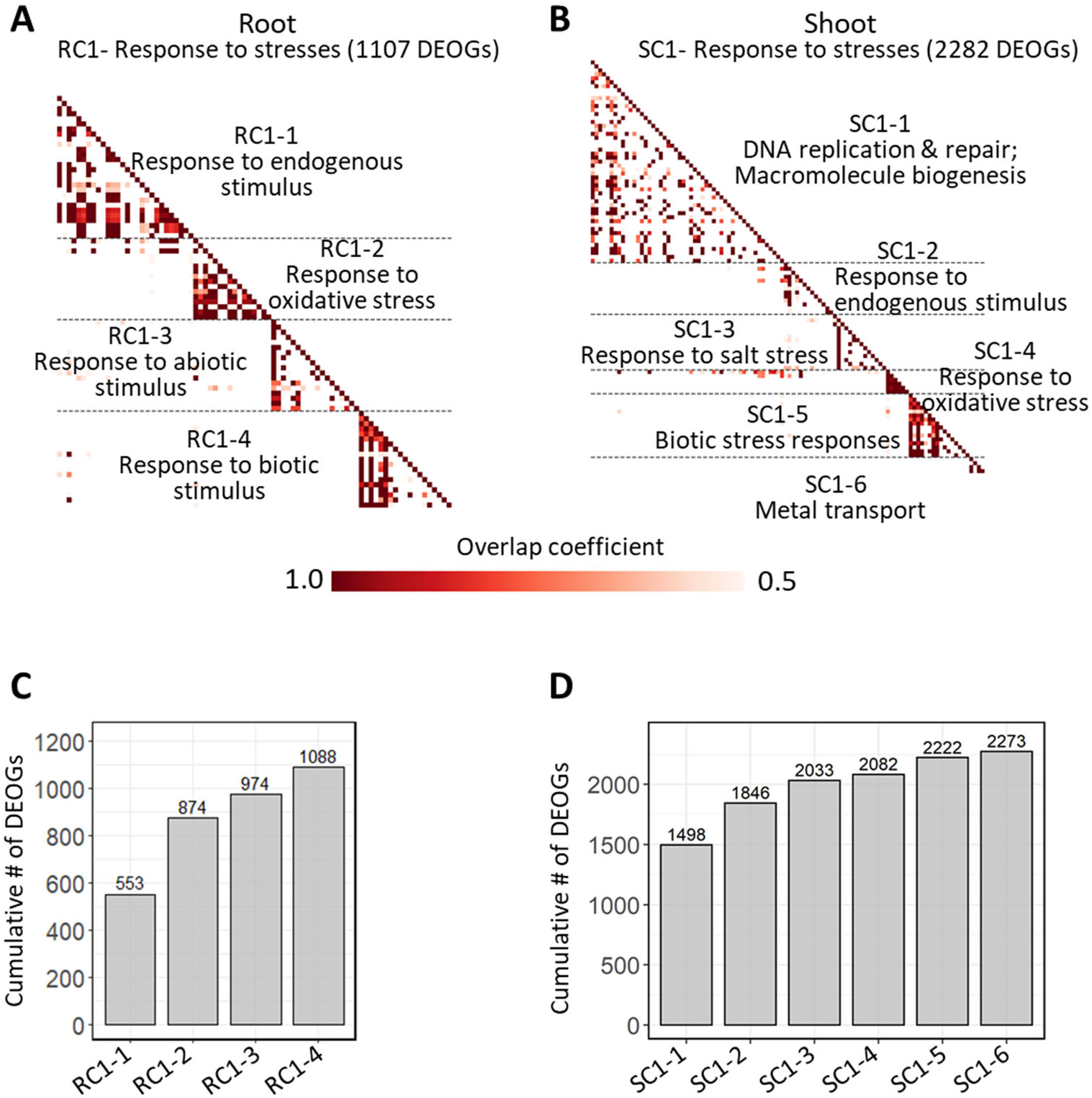
Functional processes in subclusters of the largest functional cluster (response to stresses) shown in Figure 3C. [A] Root RC1 cluster and its sub-clusters. [B] Shoot SC1 cluster and its sub-clusters. Similarity between functional clusters are based on overlap coefficient determined using GOMCL. Cumulative gene representation for sub-clusters in RC1 and SC1 for [C] roots and [D] shoots. Differentially expressed ortholog groups (DEOGs) are counted for cumulative representation in each subcluster. RC: root cluster; SC: shoot cluster.

**Figure S5.**
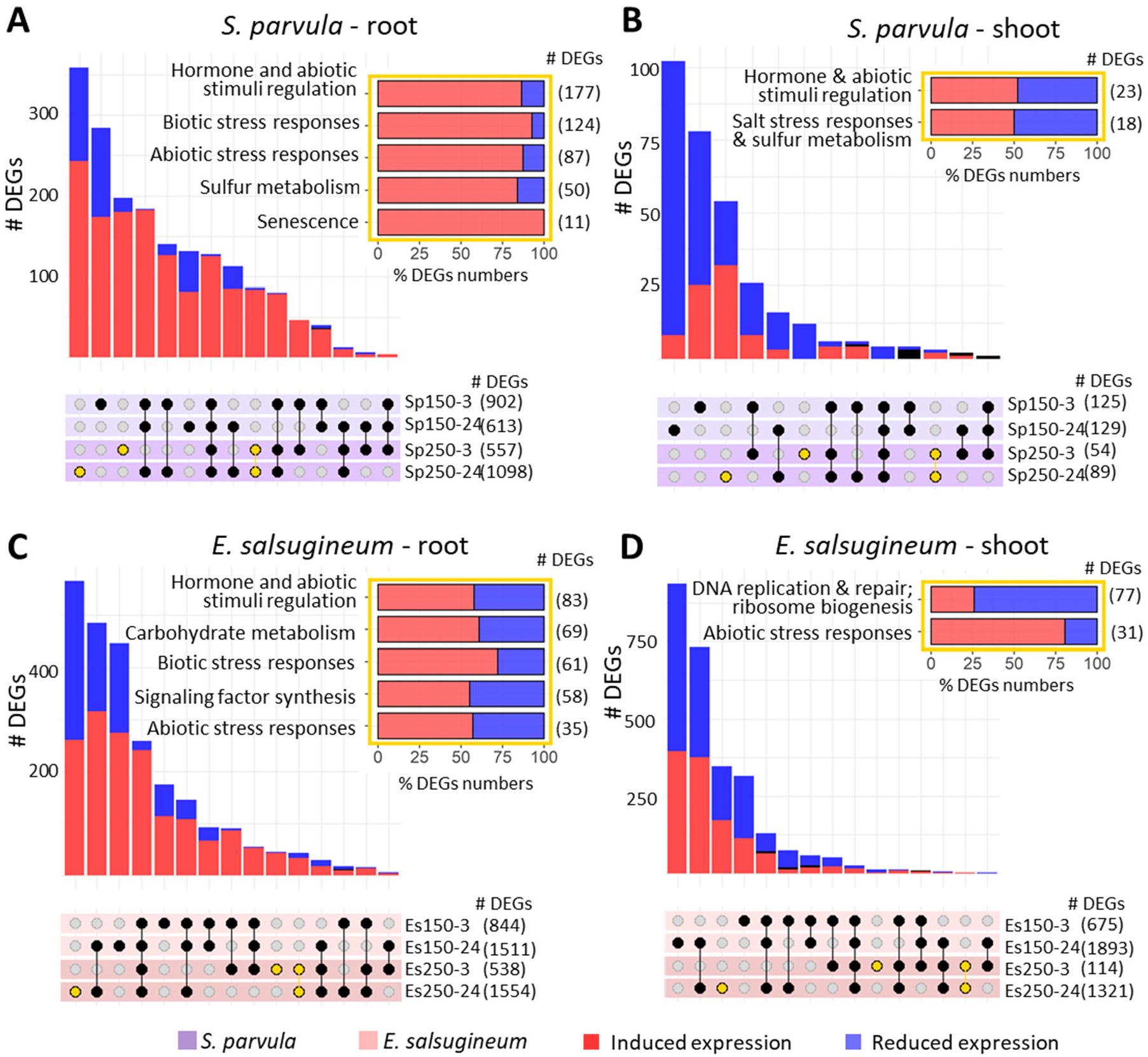
Species dependent responses to salt treatments in *Schrenkiella parvula* and *Eutrema salsugineum*. Differentially expressed genes (DEGs) in *S. parvula* [A] root and [B] shoot and *E. salsugineum* [C] root and [D] shoot in response to 150 and 250 mM NaCl stress at 3 and 24 hr. Yellow points in each panel show DEGs that were uniquely differentially expressed at 250 mM NaCl treatments. Functionally enriched processes for these unique DEGs are shown in the horizontal bar graphs given as insets outlined in yellow for each panel. Differentially expressed genes were identified at *p-*adj ≤ 0.01 (n = 3, at least 4 plants per replicate).

**Figure S6.**
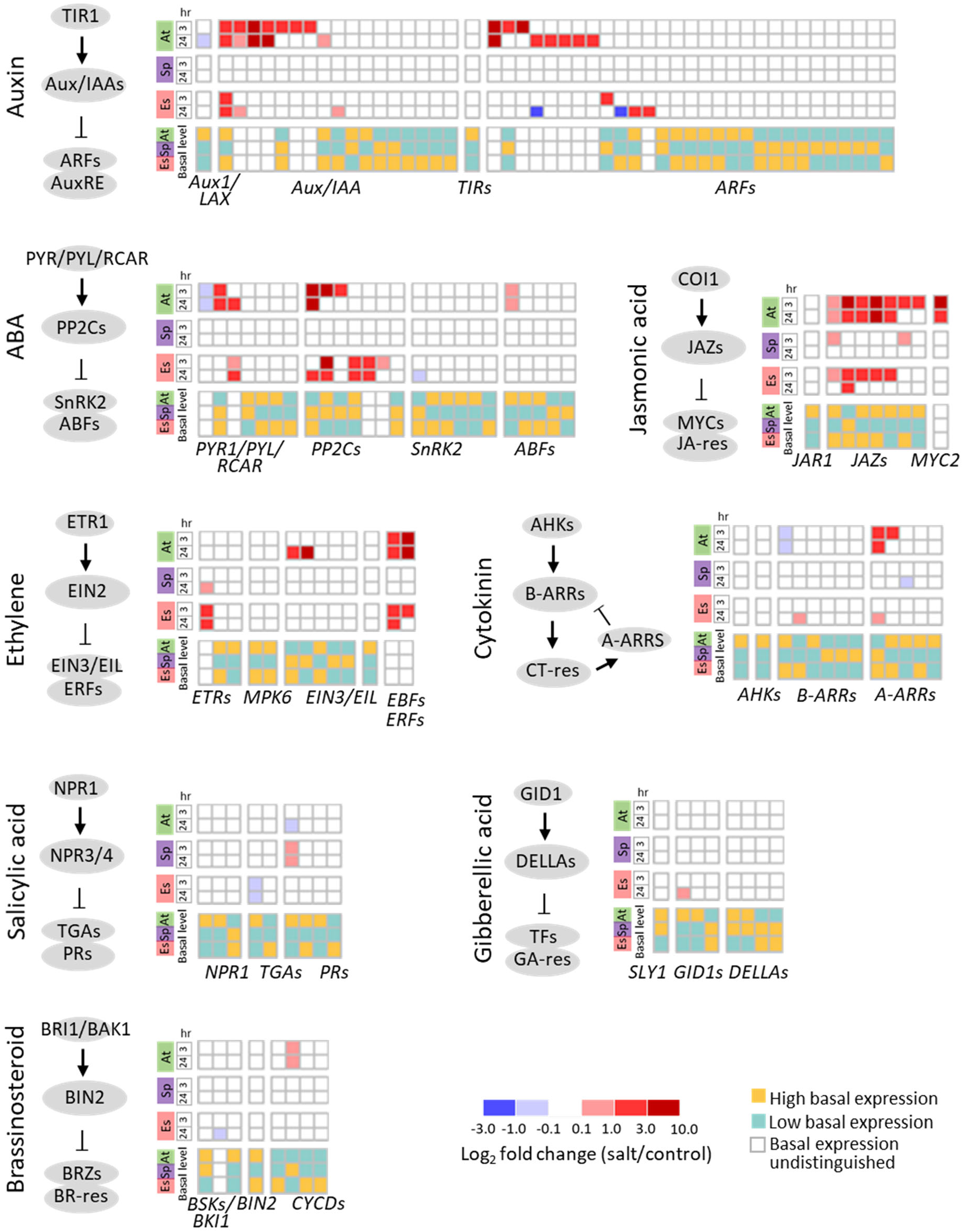
Expression of orthologs associated with hormonal signaling pathways in response to salt in *Arabidopsis thaliana* (At)*, Schrenkiella parvula* (Sp), and *Eutrema salsugineum* (Es). Genes were selected based on their KEGG annotations. Only genes that were significantly different at either basal expression among species or under salt treatments compared to the control are shown.

**Figure S7.**
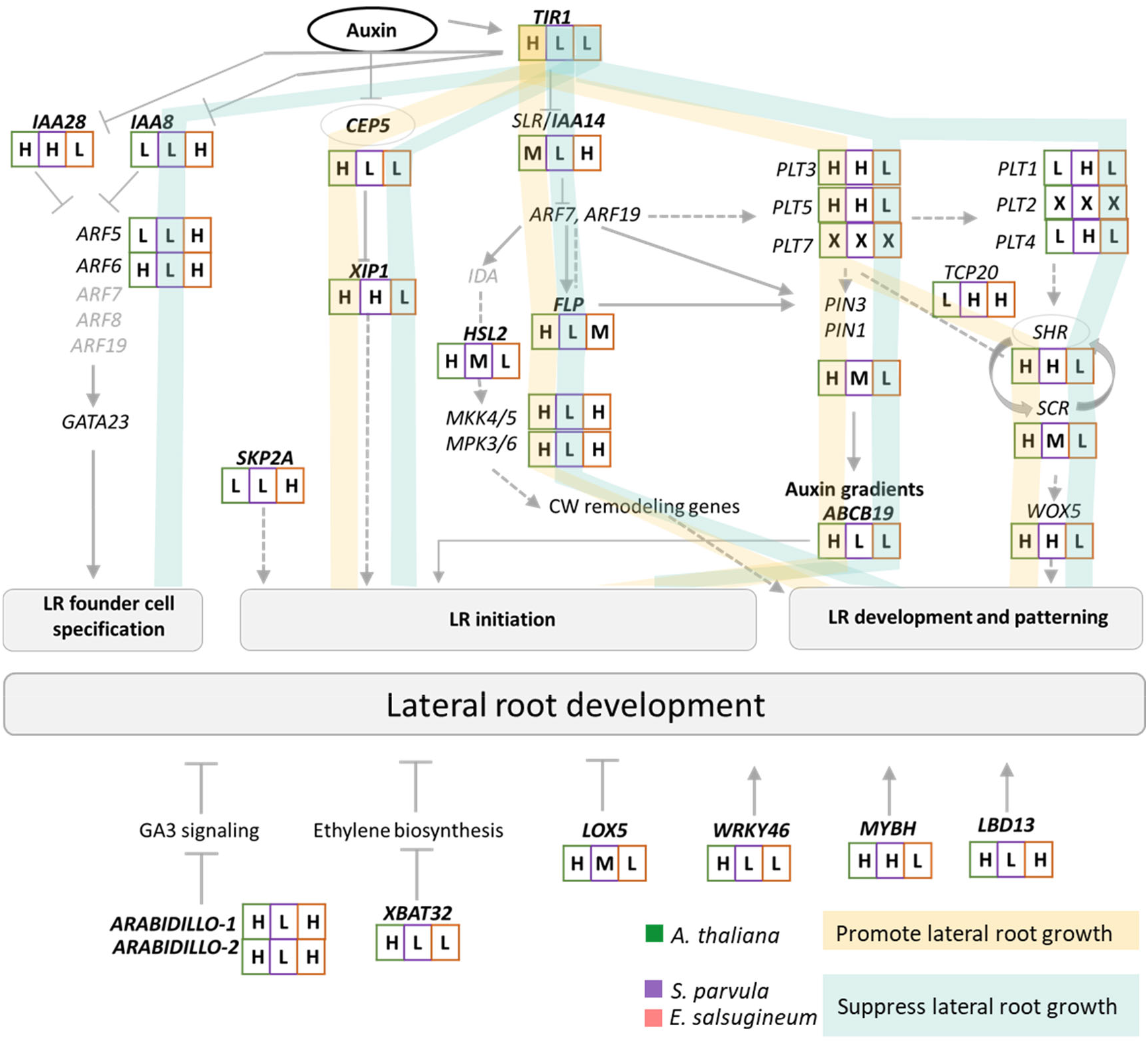
Lateral root development gene network adapted from De Rybel et al., 2010^32^ and Banda et al., 2019^33^. The expression levels of orthologs (in black) were represented as high (H), medium (M), or low (L) relative to each other at basal expression in each ortholog group. Regulatory pathways that were consistently modulated in *S. parvula* or *E. salsugineum* implied to inhibit lateral root formation are marked in turquoise ribbons. Expression pathway leading to the induction of lateral root formation in *A. thaliana* is highlighted in gold ribbons. Genes in the network that were not differently expressed among the three species are marked in light grey.

**Figure S8.**
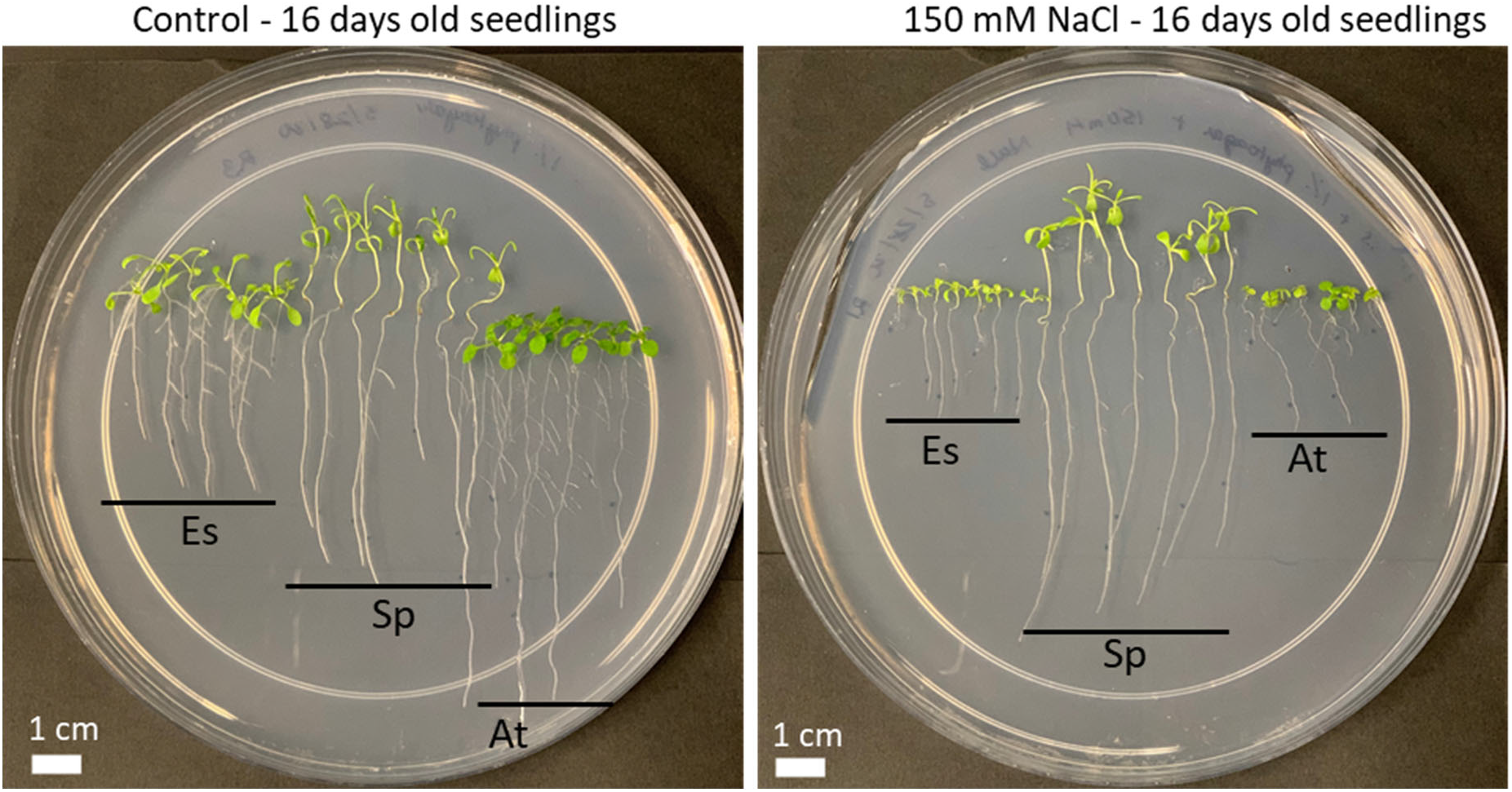
*Eutrema salsugineum*, *Schrenkiella parvula*, and *Arabidopsis thaliana* seedlings treated with 150 mM NaCl for 11 days. *Schrenkiella parvula* (Sp) promoted uninterrupted primary root growth while *E. salsugineum* (Es) showed slower primary root growth with reduced lateral root number. *A. thaliana* (At) roots showed severe primary and lateral root growth inhibition under salt stress.

**Figure S9.**
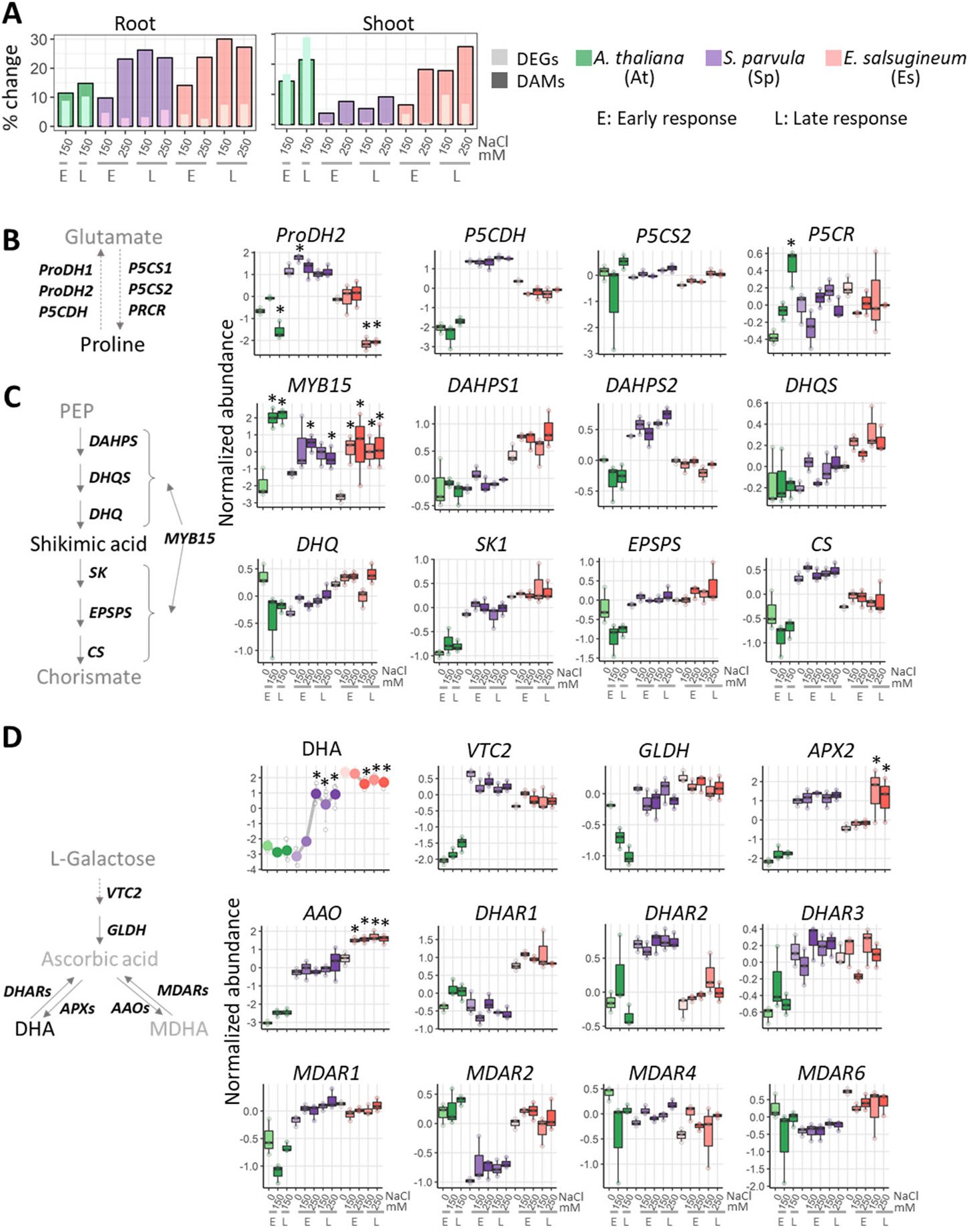
Overall concordant metabolomic and transcriptomic responses and proline, shikimic acid, and ascorbic acid pathway alignments between genes and metabolites. [A] Percentage of differentially expressed genes (DEGs) and differently abundant metabolites (DAMs) in response to salt treatments. Pathways involved in [B] proline metabolism, [C] the conversion from phosphoenolpyruvate to chorismite, and [D] ascorbic acid metabolism. Line graphs represent normalized log2 relative metabolite abundance. Boxplots represent normalized gene expression values. Center line in the boxplots indicate median; box indicates interquartile range (IQR); whiskers show 1.5 × IQR. Asterisks indicate significant difference between the treated samples and their respective controls (n = 3-4). Early (E) and late (L) responses for transcripts refer to 3 and 24 hr, respectively. E and L responses for metabolites refer to 24 and 72 hr, respectively. Metabolites are shown in the backbone of the pathway while genes encoding for key enzymes/transcription factors are placed next to the arrows. Metabolites that were quantified in the current study are given in black while those not quantified are in grey.

